# Diverse origins and transcriptional profiles of macrophages in a spinal lesion in zebrafish

**DOI:** 10.64898/2026.06.25.734502

**Authors:** Kim Heilemann, Elena Vanessa Jidav, Özge Çark, Stephen J. Enos, Bastian Wölf, Catherina G. Becker, Thomas Becker, Alberto Docampo-Seara

## Abstract

The injury responses of tissue-resident macrophages in the CNS (microglia) and blood-derived macrophages (BDMs) play key roles in successful regeneration of the zebrafish spinal cord, but the origins and dynamic behaviours of these immune cells are not well characterized. Here, we find that microglia, labelled by the *p2ry12:GFP* reporter gene, migrate long-distance through neural tissue from the brain to the spinal lesion site, while BDMs, labelled by the *mpeg1:mCherry* reporter gene, migrate mainly from the caudal hematopoietic tissue to the lesion and back. Half of *p2ry12:GFP-*positive microglia co-express *mpeg1:mCherry*, while *mpeg1:mCherry*-positive BDMs are mostly *p2ry12:GFP-*negative. However, a BDM sub-population starts to express *p2ry12:GFP* in the lesion. This indicates heterogeneous and dynamic gene expression in macrophage populations. Gene expression profiling reveals several microglia-like and BDM-like clusters in the lesion with gene expression profiles related to proliferation, phagocytosis, pro- and anti-inflammatory phenotypes and distinct expression of regeneration-relevant genes. The most abundant cell cluster are densely-packed microglia-like cells in the lesion core, which express the novel marker *g0s2*, as well as phagocytosis-related genes. Hence, regenerative success of the zebrafish spinal cord is linked to a heterogeneous and dynamic response of microglia and BDM subpopulations.

## INTRODUCTION

Spinal cord injury (SCI) causes permanent loss of motor and sensory function in mammals due to the limited regenerative capacity of the central nervous system. Following injury, a prolonged inflammatory response contributes to secondary tissue damage and promotes the formation of glial and fibrotic scars that inhibit axonal regrowth (Tran et al., 2018; Zheng & Tuszynski, 2023). In addition, neural progenitors fail to efficiently replace lost neurons and instead generate astroglia, further contributing to scar formation (Barnabé-Heider et al., 2009; Stenudd et al., 2022). In contrast, zebrafish can regenerate multiple tissues and organs, including the spinal cord (Gemberling et al., 2013).

Macrophages are key regulators of regeneration in zebrafish (Petrie et al., 2014; Klatt et al., 2026; Tsarouchas et al., 2018). Larval zebrafish are particularly suited to the analysis of the immune reaction to spinal lesion (Alper & Dorsky, 2022). At larval stages, the innate immune system, consisting mainly of neutrophils and macrophages, is the principal component of the immune system. Around 3 to 6 weeks post-fertilization, the adaptive immune system matures and becomes functional (Franza et al., 2024), such that, before that time, the functional response of the innate immune system to a spinal lesion can be studied without the influence of adaptive immunity.

Macrophages comprise both blood-derived macrophages (BDMs) and microglia, the resident macrophages of the central nervous system. These cells have different origins. BDMs derive from different sources along development, but between 3 and 5 days post-fertilization (dpf), hematopoietic stem cells that give rise to BDMs, as well BDMs itself, are located in the caudal hematopoietic tissue (CHT) (Franza et al., 2024; Jin et al., 2009; Murayama et al., 2006). In contrast, microglia originate from primitive macrophages in the rostral hematopoietic tissue (RHT) in zebrafish and colonize the brain early in development, where they properly differentiate into microglial cells (Herbomel et al., 2001; Xu et al., 2015). From ∼3 dpf on, these cells further populate the spinal cord.

In contrast to BDMs, which are continuously replenished, microglia exhibit low turnover and are maintained through local self-renewal within the CNS (Ferrero et al., 2018; Kuil et al., 2020; Shimizu & Prinz, 2025). A similar scenario is observed in humans, where microglia colonize the brain during early embryogenesis and persist as a self-renewing population largely independent of peripheral input (Réu et al., 2017). Notably, transcriptomic data have revealed a high degree of conservation between zebrafish and human microglia in both developmental dynamics and gene expression profiles (Mazzolini et al., 2020).

Following complete spinal cord transection at 3 dpf, zebrafish larvae are able to recover function within 48 hours after injury (Ohnmacht et al., 2016; Wehner et al., 2017). During this process, immune cell recruitment can be directly observed in transparent larvae (Tsarouchas et al., 2018). Neutrophils are recruited early to the injury site, peaking in number at 4-6 hours post lesion (hpl) (De Sena-Tomás et al., 2024; John et al., 2025; Tian et al., 2026; Tsarouchas et al., 2018), while BDMs and microglia are recruited to the injury site after, peaking in numbers at 24 hpl, and maintain a large presence at 48 hpl. This time course coincides with axonal regrowth, regenerative neurogenesis, and recovery of locomotion (Ohnmacht et al., 2016; Tsarouchas et al., 2018; Wehner et al., 2017). In the absence of macrophages in *irf8* mutants, increased levels of neutrophils and *il1b* expression are observed, which in turn impairs regeneration (Oprişoreanu et al., 2023; Tsarouchas et al., 2018). This highlights the importance of BDMs and microglia as key cell types in the resolution of inflammation and control of the regenerative process.

Macrophages are highly heterogeneous and can adopt diverse functional states. Macrophage activation has historically been described using the M1 (pro-inflammatory) / M2 (anti-inflammatory reparative) polarization framework in mammals (Mantovani et al., 2002, 2004). Importantly, macrophages displaying M1/M2-like polarization states have also been identified in fish (Nguyen-Chi et al., 2015; Wentzel, Janssen, et al., 2020). However, macrophage states exist along a continuum and are tailored to the specific requirements of individual tissues and regenerative contexts (Murray et al., 2015; Paolicelli et al., 2022). Consistent with this view, distinct macrophage populations with specialized functions during zebrafish spinal cord regeneration have been identified: 1) a specific subpopulation of BDMs, which promotes regenerative neurogenesis via TNF signalling (Cavone et al., 2021); and 2) a population of *sema4ab*-expressing microglia, which attenuate regenerative neurogenesis via Sema4ab-Plxnb1 and control *tgfb3* production in fibroblasts (Docampo-Seara et al., 2026).

Despite distinct roles of subpopulations of macrophages, the full complement and dynamics of different BDM and microglia populations in the spinal injury site of zebrafish has not been identified. Here we find that resident microglia in the brain proliferate and migrate to the spinal cord after spinal cord injury, while BDMs are recruited from peripheral tissues, mainly from the CHT. Overall, we identify seven BDM- and microglia-like subpopulations by their gene expression profiles across data sets, relating to proliferation, phagocytosis, and differential expression of regeneration-relevant genes. This study informs the analysis of complex immune cell interactions that facilitate spinal cord regeneration in zebrafish.

## RESULTS

In order to gain insight into the dynamics of microglia and BDMs after a spinal injury, we imaged almost the entire larvae using the double transgenic line *p2ry12:GFP;mpeg1:mCherry* in time-lapse microscopy from 3 to 33 hpl at 3 dpf (Fig. 1A). In the *p2ry12:GFP* line, the *p2ry12-*GFP fusion protein is expressed under the control of the regulatory sequences of *p2ry12* and reports microglial cell identity (Sieger et al., 2012). In the *mpeg1:mCherry* line, mCherry is expressed under the control of the regulatory sequences of *mpeg1.1*, a general marker of macrophages, including both microglia and BDMs (Ellett et al., 2011).

**Figure 1.**
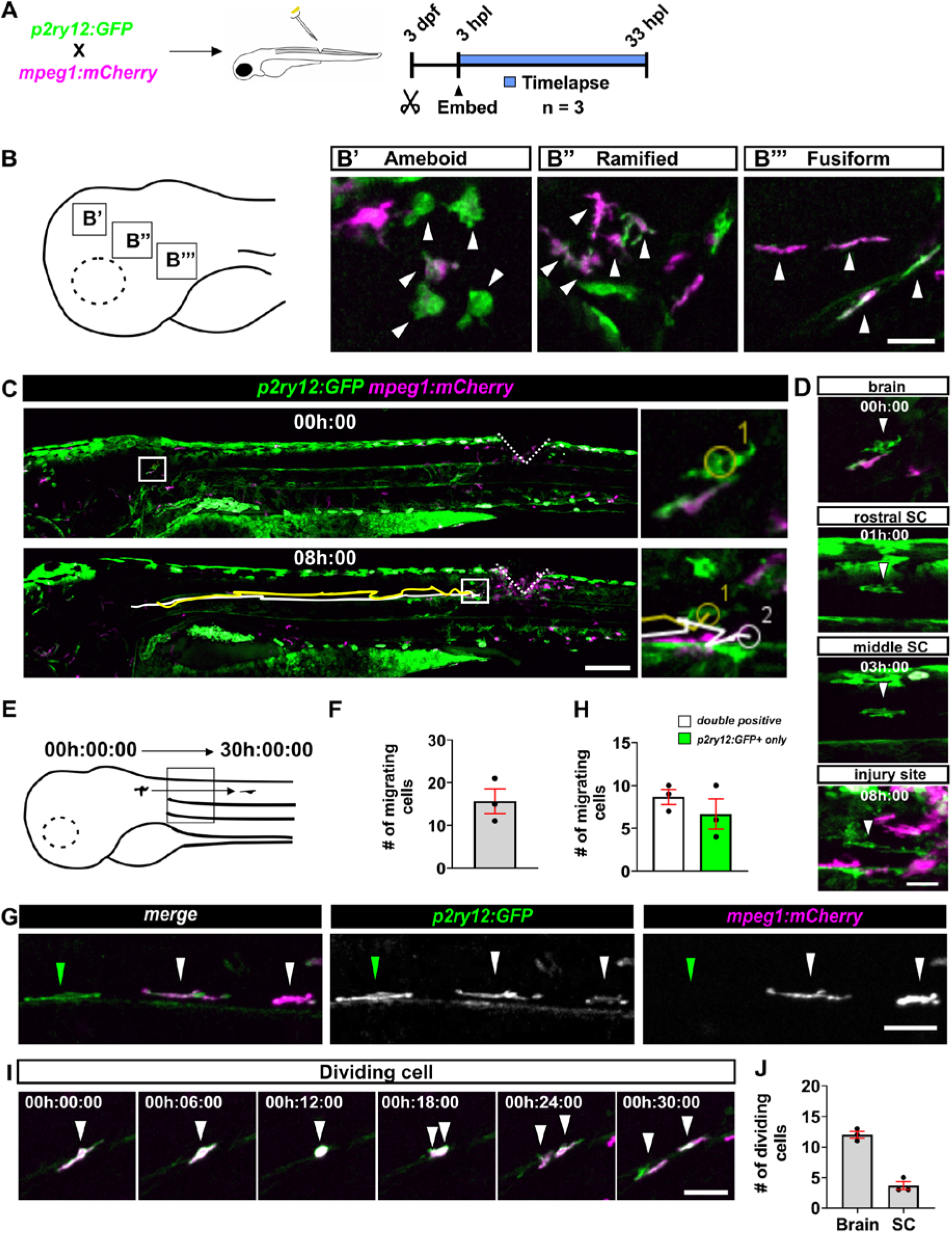
Microglia migrate towards the injury site from the brain. **(A)** Schematic and photomicrographs at different time points of the time lapse movie (0h, 1h, 3h and 8h) showing the pattern of *p2ry12:GFP* positive cells in the brain (A’, yellow arrows) and *mpeg1:mCherry* positive cells in non-neural tissue (A’’, cyan arrows). **(B)** Photomicrographs showing morphological heterogeneity (B’: amoeboid; B’’: ramified; B’’’: fusiform) of microglial cells in the brain. **(C)** Images showing the tracking over time of a single cell from the brain to the spinal cord lesion site. An average of 2-3 cells could be continuously tracked per time lapse movie. **(D)** A migrating microglial cell from the brain to the spinal cord is shown. **(E)** Schematic of how cells were quantified in F and H. We counted different reporter expression of cells coming from the brain and migrating to the spinal cord during the whole time-lapse movie in a 500 µm window placed in the brain-spinal cord boundary. **(F)** Quantification of migrating cells showing an average of 15.6 ± 2.9 cells migrating to the spinal cord. **(G)** Example images quantified in H. White arrow points at double positive cells and green arrow at *p2ry12:GFP* only positive cells. **(H)** Quantification showing that 55.3% of migrating cells were *p2ry12:GFP*-positive only, while 44.7 % shared also the *mpeg1:mCherry* signal. **(I)** Consecutive frames of a cell in the brain undergoing mitosis are shown. **(J)** Quantification of proliferating events in the brain and spinal cord. An average of 12 ± 0.5 cells were dividing in the brain and 3.7 ± 0.7 cells in the spinal cord. **Scale bars:** 500 µm (C), 15 µm (B-B’’’, D, G, I).

To determine the starting point for our analysis, we assessed transgene expression in unlesioned animals at 3, 4,and 5 dpf. Cells labelled by either transgene that were located in the CNS were considered microglia. As expected, most microglial cells were round or elongated and resided in the brain at these time points (Suppl. Fig. 1A) (Herbomel et al., 2001). These cells showed *mpeg1:mCherry* expression and low levels of *p2ry12:GFP.* Additionally, some microglial cells were detected in the spinal cord, expressing *p2ry12:GFP, mpeg1:mCherry* or both transgenes from 3-5 dpf. These cells were present in very low numbers (1.3 ± 0.63 cells per animal at 3 dpf, 3.7 ± 0.48 cells per animal at 4 dpf, 2.8 ± 1.11 cells per animal at 5 dpf; Suppl. Fig. 1B). Outside the CNS, *p2ry12:GFP*-positive cells were very rare, but *mpeg1:mCherry*-positive cells were widely distributed and particularly concentrated in the CHT (Jin et al., 2009; Murayama et al., 2006) (Suppl. Fig. 2A). Hence, unlesioned animals showed low densities of microglia or BDMs in the spinal cord to at least 5 dpf.

### Microglial cells are resident in the brain and migrate towards the injury site after spinal cord lesion

After a spinal lesion at 3 dpf, we mounted larvae to image from a lateral perspective, including head and lesion site and started time-lapse recording, after a technical delay, from 3 hpl to 33 hpl. Microglial cells with different dynamic morphologies were detected in the brain (Fig. 1B) throughout the entire time-lapse movies, including amoeboid (Fig. 1B’), ramified (Fig. 1B’’) and fusiform morphologies (Fig. 1B’’’). Fusiform cells were present relatively caudal, close to the brain-spinal cord boundary, were highly motile, and migrated towards the spinal cord. We were able to continuously track 2-3 cells from the brainstem to the lesion site (Fig. 1C-D). On average, 15.6 ± 2.9 cells per animal left the brain and invaded the spinal cord within 30 h (Fig. 1E, F). Since these numbers were much higher than the numbers of microglia present in unlesioned larvae up to 5 dpl, we concluded that this migration was injury-induced.

Of the migrating microglia in the spinal cord, 8.7 ± 0.9 cells per animal were only positive for *p2ry12:GFP*, and 6.7 ± 1.7 cells per animal were *p2ry12:GFP/mpeg1:mCherry* double-positive (Fig. 1G, H). No cells only positive for *mpeg1:mCherry* were detected. This indicated heterogeneity of microglia with respect to transgene expression.

Microglia mostly showed monotonous migration to the lesion. However, interruptions of migration occurred when microglial cells divided during migration. Cell divisions were defined by stereotypical rounding of cells with subsequent appearance of two small cells in the same place. We detected division events in the brain (12 ± 0.58 cells/time-lapse movie) and in migrating cells in the spinal cord (3.7 ± 0.67 cells/time lapse movie) (Fig. 1I, J). In summary, microglia exhibited injury-induced migration from the brain to the lesion, were heterogeneous in terms of transgene expression and showed occasional divisions.

### BDMs invade the injury site mainly from the CHT

In the same preparations as above, BDMs (*mpeg1:mCherry*-positive; located outside the CNS) migrated through non-neural tissue from multiple directions to invade the injury site. These cells acquired elongated morphologies during migration (e.g. migrating cells from CHT, arrows in Fig. 2A). Overall, 66.6% of *mpeg1:mCherry*-positive cells migrating towards the lesion came from the CHT ventral to the lesioned spinal cord, and 33.3% had other non-neural tissue origins. We noted that BDMs migrated towards and away from the lesion, mainly ventrally to rejoin the CHT. Quantifications indicated most migration toward the lesion during early time points after lesion (<12 hpl), whereas migration away from the lesion predominated at later time points (>24 hpl) (Fig. 2B). Few BDMs showed changes in the direction of migration before reaching either lesion or CHT (∼2 cells per timelapse). Interestingly, we never observed any *p2ry12:GFP-*positive cells migrating outside of the CNS. Note that at the start of the movies (3 hpl), some *mpeg1:mCherry*-positive had already reached the lesion site, consistent with previous reports (Tsarouchas et al., 2018). Overall, these observations are consistent with BDMs being mainly recruited from the CHT to the lesion and later returning to the CHT.

**Figure 2.**
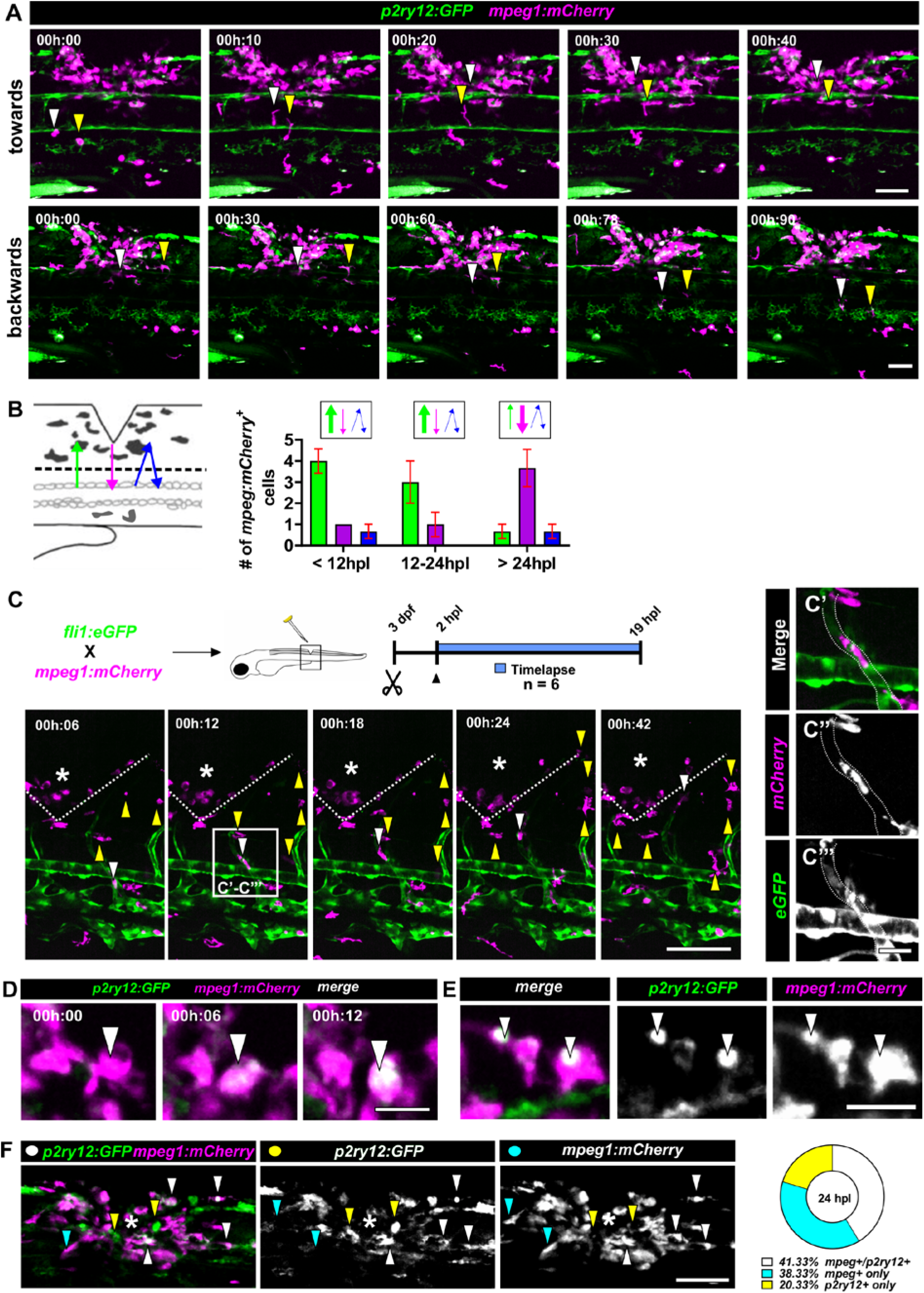
BDMs migrate to the injury site with heterogeneous routes. **(A)** *mpeg1:mCherry*-positive cells migrate towards and backwards the CHT to the injury site (arrowheads of each colour points at an individual cell over time). **(B)** Quantification of *mpeg1:mCherry*-positive cells migrating from the CHT to the injury site (green), from the injury site to CHT (magenta), and inconsistently backwards and forwards (blue) during the time lapse movies. In the first 12 to 24 hours, BDMs primarily migrate from the CHT towards the injury site, thereafter from the injury site towards the CHT. **(C)** Time-lapse movie frames showing migration to the injury site of few *mpeg1:mCherry*-positive cells through blood vessels (white arrow). The majority of the tracked cells follow different paths through the tissue (yellow arrows). **(C’-C’’’)** High magnification images showing a *mpeg1:mCherry*-positive cell inside of a blood vessel. **(D)** Timeframes showing increase of *p2ry12:GFP* signal in *mpeg1:mCherry* expressing cells. **(E)** Images showing localized subcellular presence of *p2ry12:GFP* in some *mpeg1:mCherry* positive cells. **(F)** Images and quantification of *mpeg1:mCherry, p2ry12:GFP* and double-positive cells in the injury site at 24 hpl form time lapse videos (cf. Fig. 1). White arrows point at double positive cells, yellow arrows point at *p2ry12:GFP*-positive only, and blue arrows point at *mpeg1:mCherry*-positive only cells. **Scale bars:** 100 µm (A, C), 50 µm (F), 15 µm (C’-C’’’, D, E)

In mammals, monocytes circulate through the bloodstream and extravasate into tissues in response to specific signals, where they differentiate into BDMs (Shi & Pamer, 2011; Yang et al., 2014). Therefore, BDMs may reach their targets either by migration through tissue or by entering the injury site also from blood vessels. To determine which mode of entering the lesion BDMs follow, we performed time-lapse microscopy using the transgenic line *fli1:eGFP*, in which the vasculature was labelled by expressing GFP under the control of the *fli1* promotor (Roman et al., 2002), in combination with the *mpeg1:mCherry* transgenic reporter. Time lapses run from 2 hpl to 19 hpl and were focused on the trunk at the injury site level (Fig. 2C). We were able to track 1.8 ± 0.48 *mpeg1:mCherry* positive cells per time-lapse movie inside the blood vessels, moving from the CHT to the injury area (Fig. 2C, C’-C’’’). However, the majority of BDMs (∼80% of *mpeg1:mCherry-*positive cells per time-lapse movie) were not associated with blood vessels and therefore migrated directly through different tissues to the injury site (Suppl. Fig. 2B).

Notably, we observed that some only *mpeg1:mCherry*-positive cells in the injury site showed increasing GFP signal intensity over time (Fig. 2D). Since microglial cells that only expressed *mpeg1:mCherry* and not *p2ry12:GFP* were extremely rare during migration inside the CNS, these cells likely represented BDMs that acquired microglia-like transgene expression. Quantification over the first 12 hpl, after which cells were too densely packed for reliable counts, revealed that 7.0 ± 0.58 *mpeg1:mCherry-*positive cells per animal acquired *p2ry12:GFP* expression. Of those, 3.7 ± 0.33 cells acquired diffuse cytoplasmic GFP fluorescence, suggesting upregulation of the transgene, while 3.3 ± 0.33 cells showed intense small, rounded signals, potentially related to phagocytosis of microglial debris (Fig. 2E). This indicates a scenario in which BDMs invade the injury site from surrounding tissues (mainly the CHT), where they get intermingled with microglial cells, and that at least some of these upregulate genes usually considered to be associated with a microglial identity.

### scRNA-seq analysis of macrophage-enriched cells revealed 7 subpopulations of microglia/BDMs

The above observations indicated considerable heterogeneity and dynamics of microglia and BDMs in the lesion. For example, in our time-lapse movies at 24 hpl, during active axonal and neuronal regeneration (Ohnmacht et al., 2016; Wehner et al., 2017), we observed that 41 % of the macrophages identified by either transgene at the injury site were double-positive for *mpeg1:mCherry* and *p2ry12:GFP* (microglia and BDMs with acquired microglia characteristics), 38 % were only *mpeg1:mCherry-*positive (mostly BDMs), and 20 % were *p2ry12:GFP-*positive only (microglia) (Fig. 2F), indicating considerably diversity of macrophage-related cells during active regeneration.

To investigate this diversity at higher resolution, we performed unsupervised clustering of cell profiles in a previously generated single-cell RNA sequencing (scRNA-seq) dataset of immune cells at 24 hpl (Cavone et al., 2021) (Fig. 3A). This dataset had been generated from transgenic reporter zebrafish larvae by FACS-sorting *mpeg1:GFP-*positive cells from the injury site and from comparable tissue in uninjured control fish. We subset the data for *mpeg1.1*-enriched clusters, representing BDMs and microglia (Fig. 3B). Unsupervised clustering of this subset revealed 7 different clusters (Fig. 3C). Cluster numbers 1, 3 and 4 were predominantly present after injury and therefore likely represented injury-reactive macrophage-related cells (Fig. 3D, E).

**Figure 3.**
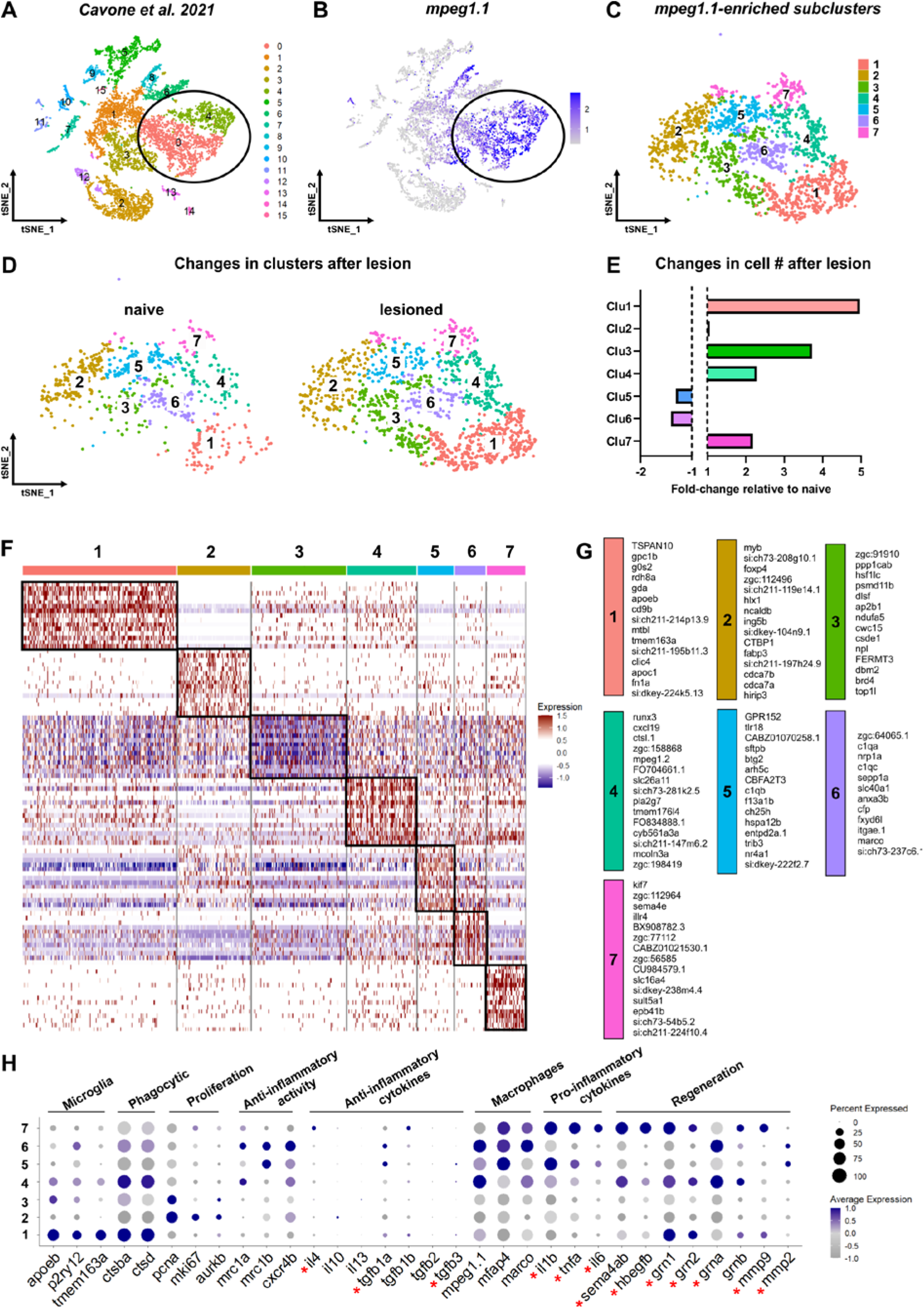
Identification of macrophage cell types by scRNA-seq analysis. **(A-B)** UMAP and Feature plot showing the clusters (0 and 4) enriched in *mpeg1.1* selected for clustering, representing macrophages & microglia (circle). **(C)** Clustering of *mpeg1.1*-enriched clusters found 7 different clusters of cells. **(D-E)** UMAPs and bar chart showing the changes in cluster numbers/proportions after injury. Increases in cell number were found in clusters 1, 3, 4 and 7. **(F)** Heat map from top-ranked gene analysis showing well differentiated clusters. **(G)** Top-ranked gene lists per cluster showing differentially expressed genes. **(H)** Dot plot showing differential expression of known marker genes for immune cell identity and major macrophage and microglia functions for identified clusters. Red asterisks indicate genes with known roles in CNS regeneration.

To characterize identity and potential functions of these clusters, we combined a top-ranked gene analysis (Fig. 3F, G) with an enrichment analysis of genes associated with known cell identities and functions (Fig. 3H). For enrichment analysis, we analyzed genes commonly associated with microglial identity, namely *apoeb, p2ry12, tmem163* (Tsai et al., 2025); macrophage identity, namely *mpeg1.1, mfap4 and marco* (Elchaninov et al., 2021; Ellett et al., 2011; Ong et al., 2020); proliferation, namely *pcna, mki67, aurkb* (Tirosh et al., 2016); phagocytic activity, namely *ctsba, ctsd* (Nakanishi, 2020); anti-inflammatory action, namely *mrc1a, mrc1b, cxcr4b,* including the pro-regenerative anti-inflammatory cytokines such as *il10, il4, il11, il13, tgfb1a, tgfb1b, tgfb2, tgfb3* (Luo et al., 2024; Nguyen-Chi et al., 2015; Yao et al., 2019); pro-inflammatory (and also regeneration-related) action, namely *il1b, tnfa, il6* (Cavone et al., 2021; Kuninaka et al., 2022; Luo et al., 2024; Nguyen-Chi et al., 2015; Tsarouchas et al., 2018; Yao et al., 2019); and additional genes expressed by immune cells with known roles in regeneration, namely *sema4ab* (Docampo-Seara et al., 2026) and *hbegfb,* (Cigliola et al., 2023), granulin (*grn1, grn2, grna and grnb* in zebrafish) (Li et al., 2013; Tsuruma et al., 2018) or mmp9 and mmp2 (Andries et al., 2021; Meschiari et al., 2018) (Fig. 3H).

Based on enrichment of expression of previously selected genes, cluster 1 was enriched in microglia (*apoeb, p2ry12, tmem163a*) and phagocytic genes (*ctsba* and *ctsd*) (Fig. 3H). The microglial genes *apoeb* and *tmem163a* were also detected in the top-ranked gene analysis for this cluster (Fig. 3G). Therefore, we tentatively identified cluster 1 as phagocytic microglia. Interestingly, this cluster is enriched in the pro-regenerative genes *grn1, grn2, grnb* and *mmp9*, suggesting a potential role in regeneration.

Cluster 2 was enriched in expression of *pcna, mki67,* and *aurkb.* Interestingly, cluster 2 showed no significant enrichment of other genes in the panel (Fig. 3H). Top ranked gene analysis for cluster 2 showed an enrichment in hematopoietic stem cell genes such as *myb* and *hlx1* (Golay et al., 1996) (Fig. 3G). Therefore, we tentatively identified cluster 2 as proliferating monocytes. Of note, no enrichment in previously published pro-regenerative genes were detected in this cluster.

Cluster 3 was enriched in microglial genes (such as *apoeb;* and moderate expression of *p2ry12)* as well as with genes associated with proliferation, as indicated by expression of *pcna* and *aurkb.* Therefore, we tentatively identified cluster 3 as proliferating microglia. Top ranked-gene analysis for cluster 3 did not identify genes that were selectively expressed in this cluster across the majority of cells (Fig. 3F). Of note, no enrichment in pro-regenerative genes were detected in this cluster.

Cluster 4 was enriched in transcripts of microglia-related genes (*apoeb, p2ry12, tmem163a),* of phagocytosis-related genes (*ctsba* and *ctsd)* and of anti-inflammatory marker genes such as *mrc1a* and *cxcr4b* (Fig. 3H). Top-ranked gene analysis also showed high enrichment in expression of the inflammatory modulator *runx3* (Lotem et al., 2017) and the fish-specific immune modulator gene *nitr2b* (Yoder et al., 2010) (Fig. 3G), suggesting a potential immune regulatory role of this cluster. Therefore, we tentatively identified cluster 4 as anti-inflammatory microglia. Interestingly, this cluster is enriched in the pro-regenerative genes *sema4ab, hbegfb*, *grn1, grn2, grna* and *grnb*, suggesting a potential role in regeneration (e.g. *sema4ab* promotes axonal regeneration and functional recovery, and controls regenerative neurogenesis (Docampo-Seara et al., 2026)).

Cluster 5 was enriched in expression of BDM-related genes (*mpeg1.1* and *mfap4), and* in pro- (*il1b* and *tnfa*) and anti-inflammatory markers (*mrc1b* and *cxcr4b*) (Fig. 3H). Top ranked gene analysis also found the expression of genes related to control of the immunity and inflammatory state, such as the immune complement gene *c1qb* and the fish specific toll-like receptor 18 (*tlr18*) (Fig. 3G). Therefore, we tentatively identified cluster 5 as hybrid BDMs, because of their mixed features of pro- and anti- inflammatory gene expression. Interestingly, this cluster is enriched in the pro-regenerative genes *tnfa, il6* and *mmp2*, suggesting a potential role in regeneration.

Cluster 6 showed enrichment of transcripts for BDM markers (*mpeg1.1, mfap4* and *marco)* and anti-inflammatory markers such as *mrc1a, mrc1b, cxcr4b*. Additionally, we found an enrichment of phagocytic markers (ctsba, ctsd). Interestingly, this cluster also showed an enriched expression of *p2ry12* (Fig. 3H). This aligns with the upregulation of *p2ry12:GFP* observed in BDMs in our time lapse movies. As well as for cluster 5, top ranked gene analysis showed expression of immune complement genes (in this case *c1qa* and *c1qc*) and the activated BDM marker *marco* (Elchaninov et al., 2021) (Fig. 3G). Therefore, we tentatively identified cluster 6 as anti-inflammatory BDMs. Interestingly, this cluster is also enriched in the pro-regenerative genes *grna, grnb*, and *mmp2*.

Finally, cluster 7 showed an enrichment of transcripts for BDM-related genes ( *mfap4* and *marco).* Top-ranked gene analysis also showed a preferential expression of genes required for development of the adaptive immune system such as *kif7* (Lau et al., 2017) and involved in immune cell infiltration such as *sult5a1* (Axelsson et al., 2012) (Fig. 3G). Additionally, this cluster showed an enrichment in pro-inflammatory cytokines (*il1b, tnfa, il6*) (Fig. 3H). Therefore, we tentatively identified cluster 7 as pro-inflammatory BDMs. Importantly, this cluster is enriched in most part of pro-regenerative genes analysed in this study (*tnfa, il6, sem4ab, hbegfb, grn1, grn2, grnb,* and *mmp9*). This pattern suggest that this population might be the pro-regenerative macrophage population define by Cavone et al., 2021.

In summary, scRNA-seq analysis identifies 7 different types or states of macrophages and microglia in lesion-site enriched tissue including distinct expression of regeneration-relevant gene.

### Robust markers genes can be identified for cell clusters

To facilitate detection of the identified cell clusters across data sets and *in situ*, we aimed establish marker genes with high discriminative power. For a systematic approach, we quantified marker performance using an Area Under the Curve (AUC) approach (Pullin & Mccarthy, 2024), which integrated both expression level and the proportion of cells expressing a given gene within a cluster. AUC scores range from 0 to 1, where 1 indicates perfect discrimination of a given cluster from all others, 0.5 corresponds to no better than random discrimination, and values approaching 0 indicate negative discrimination. Based on these scores, we classified marker performance as low (<0.5), moderate (0.5–0.7), or high (>0.7). This analysis was performed on the top 5 ranked genes (ignoring non-annotated genes) per cluster. Genes expressed in fewer than 20 % of cells within a cluster were automatically excluded by the analysis pipeline.

For cluster 1, three genes showed high discriminative power (*gpc1b, g0s2, and gda*). Among these, we selected *g0s2* due to its relatively broad expression across the cluster (60.8 % of cells) (Fig. 4A). For cluster 2, *hlx1* was the only gene with high discriminative power (Fig. 4B). As expected from the heatmap (cf. Fig. 3F), cluster 3 lacked genes with strong discriminative capacity (Fig. 4C). For cluster 4, *runx3* emerged as a strong candidate marker (Fig. 4D). Clusters 5 and 6 displayed highly similar marker profiles, preventing discrimination between these two clusters. We therefore selected *c1qa* as a shared marker to discriminate cluster 5/6 identity from the other clusters (Fig. 4E). Finally, *sult5a1* showed high discriminative power and was selected as a marker for cluster 7 (Fig. 4F).

**Figure 4.**
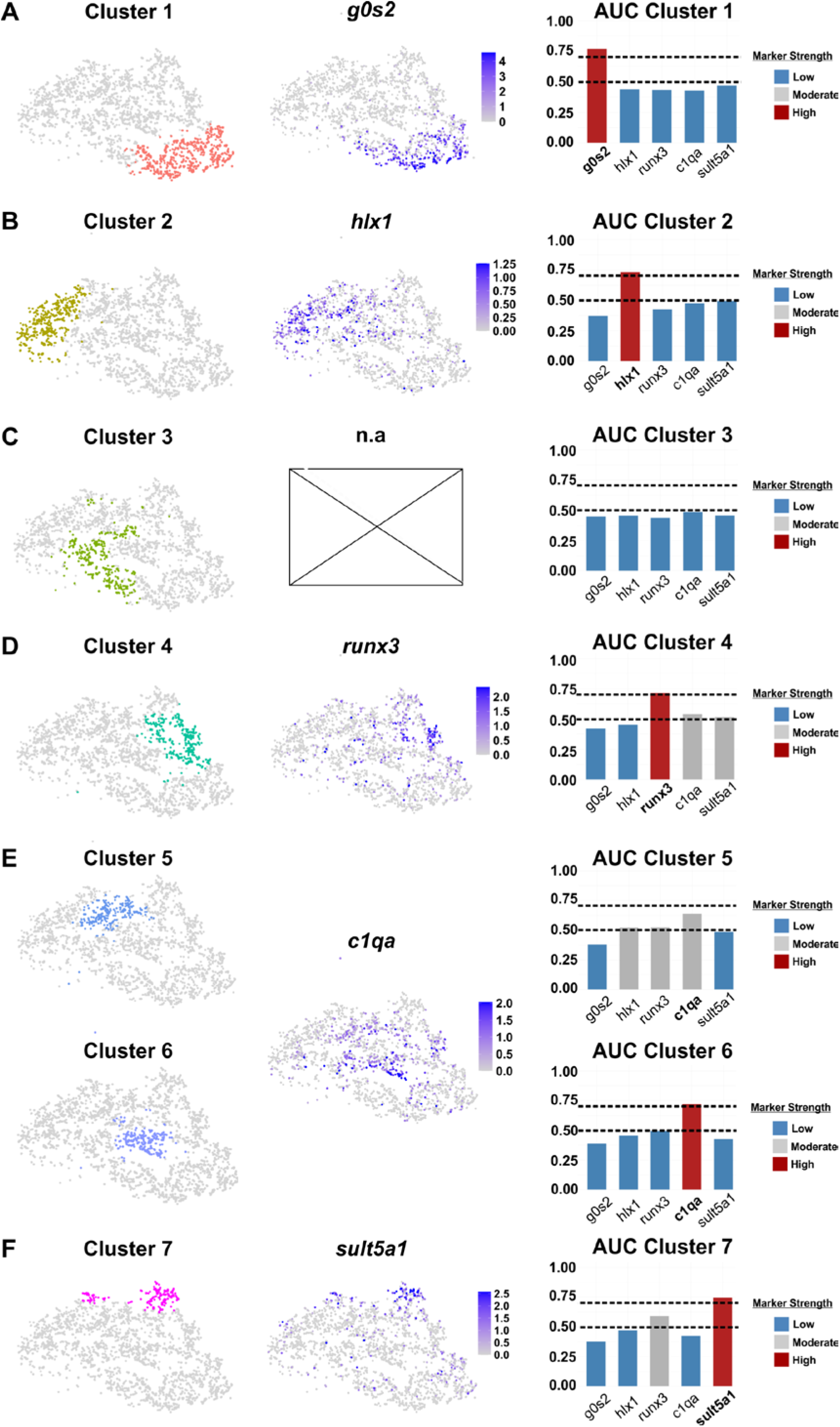
Selection of microglia/BDM clusters marker genes based on top-ranked gene expression analysis and AUC for each individual cluster. **(A)** UMAP, gene expression and AUC for cluster 1, showing enrichment and good marker strength for g0s2. **(A)** UMAP, gene expression and AUC for cluster 1, showing enrichment and good marker strength for *g0s2.* (B) UMAP, gene expression and AUC for cluster 2, showing enrichment and good marker strength for *hlx1.* (C) UMAP, gene expression and AUC for cluster 3, showing absence of good marker genes. **(D)** UMAP, gene expression and AUC for cluster 4, showing enrichment and good marker strength for *runx3.* (E) UMAP, gene expression and AUC for cluster 5 and 6, showing enrichment for *c1qa* in both clusters. **(F)** UMAP, gene expression and AUC for cluster 7, showing enrichment and good marker strength for *sult5a1.* Cluster gene is highlighted in bold in AUC analysis.

To validate these markers, we tested their expression in a previously obtained scRNA-seq data set at 24 hpl from whole lesioned tissue, i.e. without cell type enrichment by FACS (Docampo-Seara et al., 2026). In that data set, which contained all major cell types, four clusters were originally identified, with two microglia-like ones (MM#1, MM#2) and two BDM-like ones (MM#3, MM#4) (Suppl. Fig. 3A). All of our chosen markers labelled subpopulations of clusters MM#1-4 (Suppl. Fig. 3B). Microglial markers *g0s2,* labelled sub-populations of the previously identified microglial clusters. This marker was also highly selective for immune cells. *C1qa* expression was also selective for macrophages. *Runx3,* additionally to macrophages, labelled neutrophils and xantophores. *sult5a1* labelled neutrophils in addition to macrophages, and *hlx1* was also found in fibroblasts (Suppl. Fig. 3C). Thus our markers identify (sub-)types of macrophages also in whole tissue and some markers are highly selective for microglia. Hence, this analysis extends the previous analysis of macrophage subtypes in spinal cord lesion.

### Identified groups of microglia and BDMs are present *in situ* and occupy different positions in the injury site environment

To analyse the abundance and tissue distribution of the microglia and BDM clusters identified by scRNA-seq, we performed HCR-FISH for the selected marker genes at 24 hpl using the *mpeg1:mCherry* line. For each cluster, we described: 1) detection of marker expression by HCR, 2) qualitative spatial distribution within the lesion, 3) quantitative spatial enrichment using Kernel Density Enrichment (KDE) analysis (Silverman, 1998), and 4) comparison between *in situ* cell counts (proportion of *mpeg1:mCherry* positive cells expressing a marker gene) and scRNA-seq predicted proportions for each corresponding cluster. This approach allowed us to assess both the existence and spatial organization of each cluster *in situ*. For spatial distribution we used a classical body axis reference (rostra vs caudal, and dorsal vs ventral), with an addition description of the relative position respect to the injury site (peripheral to the lesion in intact tissue, in the interphase of the lesioned tissue, or in the lesion core)

For cluster 1 (phagocytic microglia), HCR-FISH confirmed abundant *g0s2/mpeg1:mCherry* double-positive cells at the injury site (Fig. 5A; Suppl. Fig. 4A). These cells were mainly distributed in the interphase of the lesioned tissue and in the lesion core. KDE spatial enrichment analysis showed strong enrichment in these areas. Interestingly, in both the scRNA-seq data and in HCR quantifications, this cluster represented the most abundant cluster.

**Figure 5.**
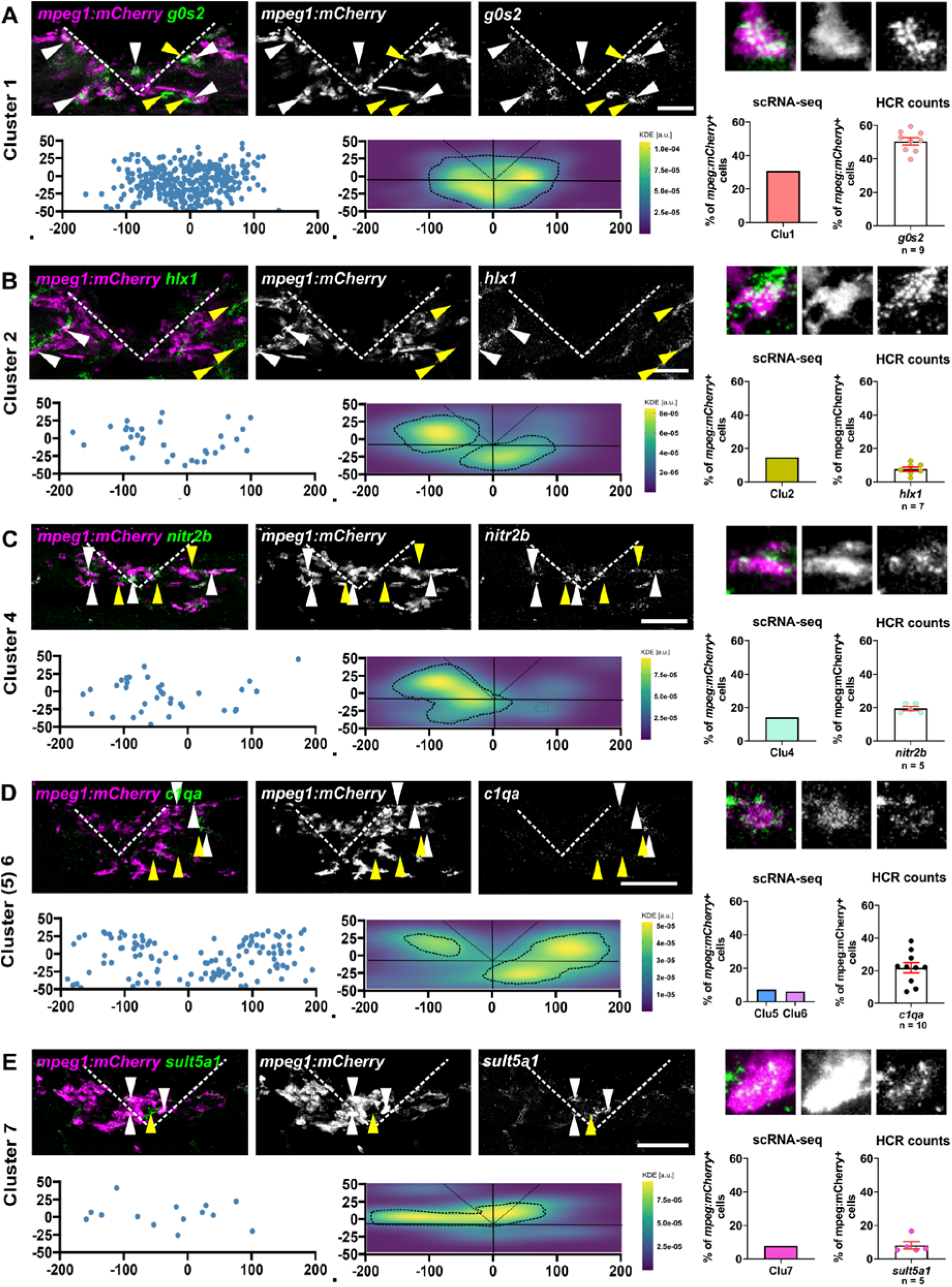
*in situ* validation and spatial characterization of selected genes by HCR in *mpeg1:mCherry* reporter larvae. **(A)** HCR, spatial distribution, KDE analysis and quantification of cluster 1 by HCR against *g0s2* in the *mpeg1:mCherry* transgenic line. *g0s2/mpeg1:mCherry* double-positive cells are pointed with white arrows. Cells positive for *g0s2*, but *mpeg1:mCherry*-negative, are pointed by yellow arrows. Double-positive cells are located in the lesion core and interphase tissue. scRNA-seq predicted 31% of cells for this cluster and in *situ* counts retrieve a value of 50%. **(B)** HCR, spatial distribution, KDE analysis and quantification of cluster 2 by HCR against *hlx1* in the *mpeg1:mCherry* transgenic line. *hlx1/mpeg1:mCherry* double-positive cells are pointed with white arrows. Cells positive for *hlx1* that are *mpeg1:mCherry*-negative are indicated by yellow arrows. Double positive cells are located peripheral to the injury site and preferentially enriched in the rostral site and ventral to the injury. scRNA-seq predicted 15% of cells for this cluster and in *situ* counts retrieve a value of 10%. **(C)** HCR, spatial distribution, KDE analysis and quantification of cluster 4 by HCR against *nitr2b* in the *mpeg1:mCherry* transgenic. *nitr2b/mpeg1:mCherry* double positive cells are pointed with white arrows*. nitr2b* positive cells that are *mpeg1:mCherry* negative are indicated by yellow arrows. Double positive cells are located close to the injury site and preferentially enriched in the rostral site in the interphase and peripheral to the injury. scRNA-seq predicted 14% of cells for this cluster and in *situ* counts retrieve a value of 20%. **(D)** HCR, spatial distribution, KDE analysis and quantification of clusters 5-6 by HCR against *c1qa* in the *mpeg1:mCherry* transgenic line. *c1qa/mpeg1:mCherry* double positive cells are pointed with white arrows. Some *c1qa* positive cells that are *mpeg1:mCherry* negative are indicated by yellow arrows. Double positive cells are located peripheral to the injury site and preferentially enriched in the caudal site scRNA-seq predicted 6% and 7% of cells for each cluster and in *situ* counts retrieve a value of 23%. **(E)** HCR, spatial distribution, KDE analysis and quantification of cluster 7 by HCR against *sult5a1* in the *mpeg1:mCherry* transgenic line. *sult5a1/mpeg1:mCherry* double positive cells are pointed with white arrows. *sult5a1* positive but *mpeg1:mCherry* negative cells are indicated by yellow arrows. Double positive cells are located in the lesion core and rostral to the injury site. scRNA-seq predicted 8% of cells for this cluster and in *situ* counts retrieve a value of 9%. **Scale bars:** 50 µm.

Several *g0s2* positive cells that were not *mpeg1:mCherry* were found by HCR. These cells could be additional microglial cells, as we observed microglial cells (*p2ry12:GFP-*positive) in our time lapse that did also not express *mpeg1:mCherry* (see Fig. 2F). To characterized these additional *g0s2*-positive cells, we performed multiplexed HCR-FISH at 24 hpl of an established microglial marker, *apoeb* (Krasemann et al., 2017), in combination with *g0s2* (Suppl. Fig. 4B). We found that 88% of *g0s2*-positive cells colocalized with *apoeb* signal. This contrasts with the number of *g0s2*/*mpeg1:mCherry* double positive cells, which represented 71% of the total *g0s2-*positive counts (see Suppl. Fig. 4A), suggesting that *g0s2* also labels a population of microglial cells which are not *mpeg1:mCherry*-positive. Additionally, we tested immunolabelling with the 4C4 microglial marker antibody, since previous studies also show that 4C4 antibody is a useful tool to identify microglia in adult zebrafish and larvae (Becker & Becker, 2001; Docampo-Seara et al., 2026; Rovira et al., 2023; Tsarouchas et al., 2018) (Suppl. Fig. 4C). We found that 75.8 % of *g0s2*-positive were also positive for 4C4. Then, we checked for *g0s2* expressing cells in the microglia deficient line *csf1ra/b* -/- (Ferrero et al., 2021; Oosterhof et al., 2018), and found a reduction of ∼80% in the number of *g0s2*-positive cells (Suppl. Fig. 4D). This suggests *g0s2* labels mostly microglial cells.

For cluster 2 (proliferating monocytes), HCR-FISH identified low numbers of *hlx1/mpeg1:mCherry* double-positive cells (Fig. 5B). These cells were located peripheral to the lesion in intact tissue. KDE spatial enrichment analysis revealed enrichment in these areas at rostral and ventral positions respect to the injury. The proportion of cluster 2 cells *in situ* was comparable to that in the scRNA-seq data. *hlx1*-positive cells that were *mpeg1:mCherry-*negative could be found outside the injury site aligned with blood vessel distribution resembling the pattern of perivascular fibroblasts (yellow arrows). Expression of *hlx1* in fibroblasts is in line with the prediction from non-FACS-sorted scRNA-seq (cf. Suppl. Fig. 3C).

For cluster 4 (anti-inflammatory microglia), the *runx3* probe did not yield detectable signal for technical reasons. To still provide a comprehensive characterization of cluster 4, we manually expanded our analysis for cluster 4 to the top 30 ranked genes, and we manually selected *nitr2b* (Yoder et al., 2010), which was partially expressed by cluster 1, but preferentially expressed in cluster 4 (44% of cells in cluster 4 vs 27% of cells in cluster 1). HCR-FISH detected *nitr2b/mpeg1:mCherry* double-positive cells distributed around the injury site (Fig. 5C). These cells were mainly distributed in the interphase of the lesioned tissue. KDE spatial enrichment analysis indicated enrichment in these areas. *In situ* quantification roughly matched scRNA-seq predictions. Number of *nitr2b*-positive cells labelling outside *mpeg1:mCherry*-positive cells was negligible, matching with the non-FACS-sorted scRNA-seq prediction.

For clusters 5 and 6 (hybrid and anti-inflammatory BDMs respectively), HCR-FISH revealed several *c1qa/mpeg1:mCherry* double-positive cells, mainly located peripheral to the lesion in intact tissue (Fig. 5D). KDE spatial enrichment analysis showed enrichment in ventro-caudal regions peripherally to the lesion in intact tissue. In contrast to other quantifications, i*n situ* counts exceeded scRNA-seq predictions, even combining both clusters 5 (6 %) and 6 (7%) together (23 % vs 13% predicted by scRNA-seq for both clusters). *c1qa*-positive cells labelling outside *mpeg1:mCherry*-positive cells was negligible, matching the non-FACS-sorted scRNA-seq prediction.

Finally, for cluster 7 (proinflammatory BDMs), HCR-FISH detected a low number of *sult5a1/mpeg1:mCherry* double-positive cells, spread in the injury site (Fig. 5E). KDE spatial enrichment analysis revealed enrichment in regions adjacent to the lesion core, particularly in dorso-rostral areas, with extension toward peripheral rostral regions. *In situ* counts closely matched the low abundance scRNA-seq predictions.

Several *sult5a1*-positive cells that were not *mpeg1:mCherry*-positive were found by HCR. Non-FACS-sorted scRNA-seq analysis predicted that these cells might be neutrophils. To validate this finding, we performed HCR against *sult5a1* in a double transgenic line reporting macrophages (*mpeg1:mCherry*) and neutrophils (*mpx:GFP*) (Suppl. Fig. 5A). We found that 72.4% of *sult5a1*-positive cells were also *mpx:GFP*-positive and only 13.3 % were positive for *mpeg1:mCherry* (Suppl. Fig. 5B). A remaining 14.3% of *sult5a1*-positive cells did not co-express any transgene. Almost half of the *mpx:GFP*-positive cells co-expressed *sult5a1* (Suppl. Fig. 5C). This suggests that, despite labelling a discrete cluster of BDMs, *sult5a1* also labels a large number of neutrophils.

Finally, to determine whether the populations defined by our marker genes shared similar spatial organizations or not, and therefore might be involved in similar/different biological processes, we performed a spatial correlation of KDE-enriched areas. The scores ranged from −1 (negative correlation or non-overlapping probability) to +1 (positive correlation or fully overlapping probability). In general, correlations between clusters were close to 0 (no better than random), indicating independent spatial distributions (Suppl. Fig. 6). This suggested that the different microglial and BDM clusters occupy segregated spatial territories within the lesion, that might be linked to their inferred function or cell dynamics. For example, phagocytic microglial cells were enriched in the edges of the site tissue, where debris was phagocytosed.

In summary, we identified 7 populations of macrophage-related cell clusters with distinct gene expression profiles across data sets and *in situ* that may have distinct function in regeneration. A summary of the spatial distribution and gene profile for the different clusters can be seen in Fig. 6.

**Figure 6.**
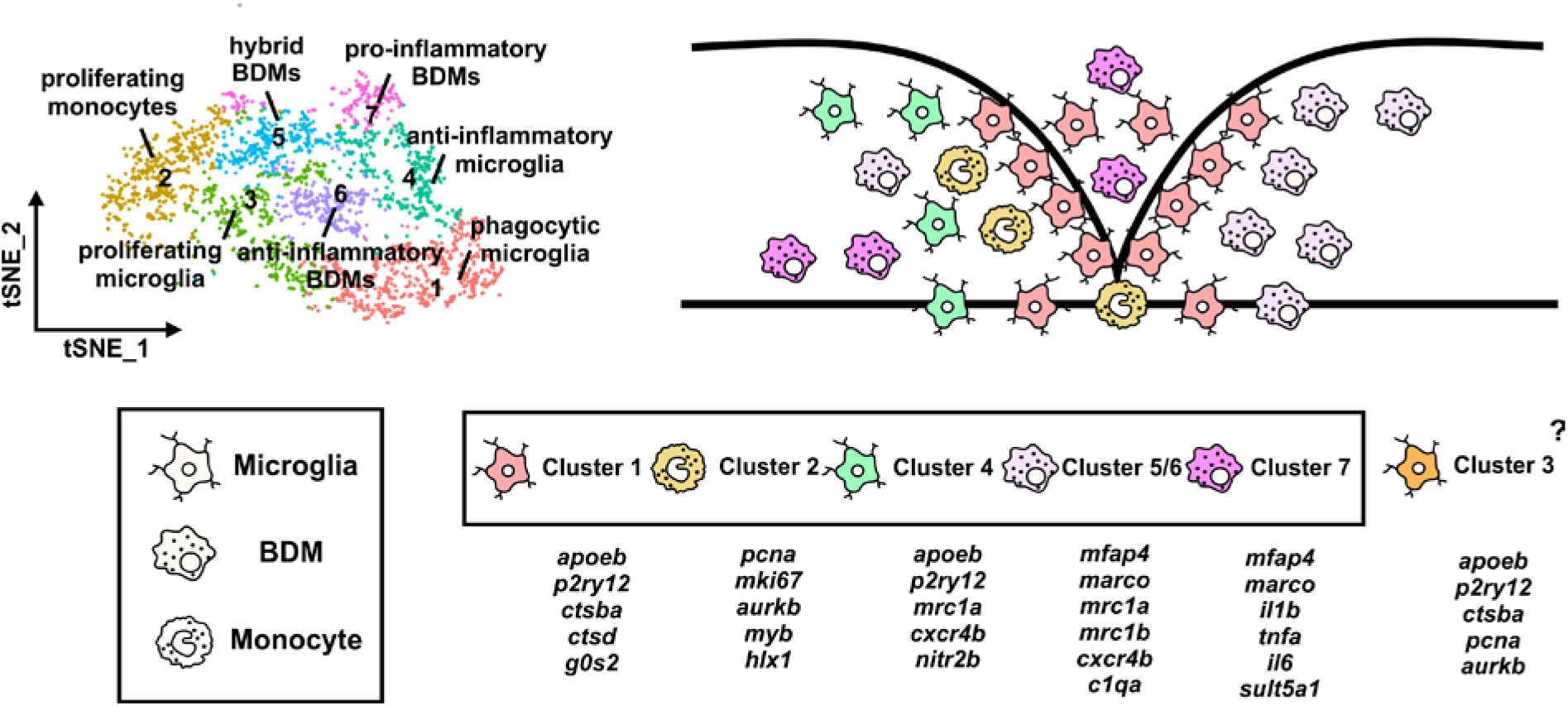
Summary of microglia/BDMs cell clusters and their distribution in the injury site. UMAP shows the identified clusters with their annotated names. Drawing of the injury site shows the different populations with their preferential locations in the lesion site. Legend shows cluster names with their enriched genes. Cluster 3 was not included in the drawing, since we could not find good substantial differential expressed genes for *in situ* validation.

## DISCUSSION

In this study, we provide a comprehensive characterization of macrophage populations during spinal cord regeneration in zebrafish larvae by integrating *in vivo* imaging, single-cell transcriptomic, and spatial validation. Our findings reveal that successful regeneration is associated with a highly dynamic and heterogeneous immune response. Microglial cells, located in the brain, travel long distances to arrive to the injury site environment, while BDMs migrate locally form the CHT and peripheral tissues; yet converge within the lesion environment to form a coordinated response. Interestingly, some BDMs, once arrived at the injury site, acquire *p2ry12* expression, normally only expressed by CNS tissue-resident microglia. scRNA-seq analysis and *in situ* validation indicated the existence of 7 different clusters of macrophages in the injury site with different transcriptional states and expression of previously identified regeneration-relevant genes.

### Microglial cells migrate from the brain after injury and show different transcriptional states that promote regeneration

Here we show that microglial cells in the brain exhibit classical ameboid and ramified morphologies previously identified in mammals in health and disease (reviewed by Savage et al., 2019) and in unlesioned zebrafish larvae (Herbomel et al., 2001), also now in injured fish. After lesion, these cells acquire a fusiform morphology associated with migration long-distance towards the injury site. Recruitment of microglial cells toward the distant injury site likely depends on long-range signalling mechanisms. Previous work in zebrafish demonstrated that neuronal damage induces propagating intercellular Ca²⁺ waves that spatially define microglial recruitment and migration toward the lesion (Sieger et al., 2012). These responses are associated with ATP/purinergic signalling through receptors such as P2ry12, a key mediator of microglial chemotaxis toward tissue damage, while glutamate-dependent signalling further contributes to the propagation of injury-induced Ca²⁺ activity (Ohsawa & Kohsaka, 2011; Sieger et al., 2012). Together, these observations support the idea that microglial mobilization after spinal cord injury is coordinated by diffusible and activity-dependent signals capable of transmitting damage information across large regions of the central nervous system.

Previous studies generated a FACS-sorted scRNAseq dataset of *mpeg1:GFP*-positive cells to provide insight into the roles of macrophages in spinal cord regeneration in zebrafish larvae (Cavone et al., 2021). The existence of a macrophage-enriched dataset allowed us to perform high-resolution analysis. By unsupervised sub-clustering of macrophages, including BDMs and microglia, we identified at least three distinct microglial populations enriched for *apoeb, p2ry12* and *tmem163*, characterized by phagocytic capacity (high cathepsin family gene expression), proliferative activity (enrichment in *pcna, mki67, aurkb*) or anti-inflammatory features (*mrc1a* and *cxcr4b* expression). Remarkably, the phagocytic and anti-inflammatory clusters were enriched in genes with functional importance in spinal cord regeneration, such as *sema4ab* and *hbegfb,* (Cigliola et al., 2023; Docampo-Seara et al., 2026) or granulins (*grn1, grn2, grna and grnb* in zebrafish) (Li et al., 2013; Tsuruma et al., 2018), supporting a functional role for these cell populations in controlling the regeneration success. These populations occupy different spatial positions within the injury site. Anti-inflammatory microglia (defined by *nitr2b* expression) are preferentially localized to rostral regions, consistent with their origin and migration path, while phagocytic microglia (defined by *g0s2* expression) accumulate at lesion tissue edge, where debris clearance takes place. Hence, spinal cord injury induces the coordinated mobilization from the brain to the injury site of transcriptionally and spatially distinct microglial populations that likely contribute to successful regeneration through specialized inflammatory and phagocytic functions.

### BDMs preferentially migrate from the CHT and represent pro- and anti-inflammatory profiles

We show that BDMs migrate to the injury site from peripheral tissues, particularly the CHT (Jin et al., 2009; Murayama et al., 2006) and infiltrate the lesion via multiple routes. While a subset of cells migrates through the vasculature, the majority arrives through surrounding tissues, indicating diverse migratory strategies. Once within the lesion site, BDMs intermingle with microglial cells.

Our scRNA-seq analysis identified 3 BDM clusters (enriched for *marco* and *mfap4*). Cytokine profile analysis shows that cluster 7 is characterized by pro-inflammatory cytokines (*il1b, tnfa, il6*), cluster 6 by anti-inflammatory markers (*mrc1a, mrc1b, cxcr4b*), and cluster 5 by a mix of both. Interestingly, this hybrid cluster had a moderate expression of *p2ry12,* but lacks of other microglial markers, such as *apoeb* or *tmem163*. This is in line with our time lapse observations where we observed a small fraction of *mpeg1:mCherry* positive cells acquiring *p2ry12:GFP* expression upon invading the spinal cord injury site. Similarly, previous studies have also found that infiltrating BDMs in the brain can acquire *p2ry12* expression in a tumour model in zebrafish (Chia et al., 2018). These findings suggest that infiltrating BDMs can partially adopt microglial-like phenotypes within the injured CNS, highlighting a remarkable degree of transcriptional plasticity in zebrafish macrophages.

In mammals, monocytes circulate through the bloodstream and extravasate into tissues in response to specific signals (Shi & Pamer, 2011; Yang et al., 2014). Consistent with these findings, our time-lapse imaging revealed a small number of blood-circulating or extravasating cells associated with blood vessels. From our scRNA-seq analysis, one of the seven clusters (cluster 2) lacks classical BDM or microglial markers, but expresses proliferation-associated genes such as *pcna* and *aurkb*, consistent with a proliferating monocyte identity. This cluster was detected at low abundance in both scRNA-seq and HCR analyses (<10%), align with low numbers detected of blood-circulating *mpeg1*-positive cells in time lapses. Taken together, scRNA-seq, HCR quantification, and time-lapse observations supports the annotation of cluster 2 as proliferating monocytes.

In summary, these findings suggest that BDMs display remarkable migratory and transcriptional plasticity after spinal cord injury, transitioning through distinct inflammatory states and partially acquiring microglial-like features within the regenerating CNS.

### Relevance of identified cell clusters for regeneration

Unsupervised clustering of immune cells in the lesion showed clusters related to different microglia and BDM gene expression profiles. Importantly, this analysis also showed regeneration-relevant gene expression also segregated between these clusters. For example, *tnfa* which promotes axon regeneration and neurogenesis (Cavone et al., 021; Tsarouchas et al., 2018), was mainly expressed in BDM cluster 7 (pro-inflammatory BDMs) and *sema4ab*, which promotes axon growth and attenuates neurogenesis (Docampo-Seara et al., 2026) was mainly expressed in that cluster and microglia cluster 4 (anti-inflammatory microglia). Hence, the identified cell clusters, representing different cell types or states likely have distinct functions in spinal cord regeneration and are therefore relevant grouping. Future analyses will have to determine how these cell populations change during the previously described a biphasic inflammatory response after zebrafish spinal cord injury, characterized by a transition from a pro-inflammatory to an anti-inflammatory cytokine abundance around 24 hpl (Tsarouchas et al., 2018).

### *g0s2* identifies a major subset of microglia

Recent studies analysing microglial cells in the context of retinal degeneration in zebrafish at a computational level found *g0s2* among the top expressed (Ravishankar et al., 2026). However, *in situ* analysis of these gene and its potential as microglial marker gene has never been tested. Here we show using scRNA-seq analysis and *in situ* validation, that *g0s2* labels the main cluster of microglial cells after spinal cord injury in zebrafish. Analyses using microglial markers *apoeb* and 4C4 showed that ∼80% of *g0s2*-positive cells co-expressed microglial marker. Moreover, the ∼80% reduction in the number of *g0s2*-positive cells in the *csf1ra/b* double mutant microglia-deficient line firmly establishes the gene as a microglia marker. Interestingly, co-labelling of *g0s2*-positive with microglia markers was more extensive than with *mpeg1:mCherry* (71%), suggesting that *g0s2*-positive that are *mpeg1:mCherry-*negative cells were potentially related to the *p2ry12:GFP*-positive/ *mpeg1:mCherry-negative* microglia cells observed in time-lapse movies. Overall this suggests that *g0s2* constitutes a reliable microglia marker after spinal cord injury in zebrafish larvae.

### Limitations of the study

Several limitations of this study should be considered. First, macrophage states were defined based on transcriptional profiles and marker expression, which do not fully resolve lineage relationships or functional roles. In particular, whether BDMs adopt stable microglial identities remains unclear and will require lineage tracing approaches. Second, overlapping gene expression between clusters limits marker specificity, necessitating combinatorial strategies for precise cell identification. Finally, computational analysis using *mpeg1:GFP* to analyse macrophage subclusters excluded *mpeg1:GFP*-negative microglia, which may have introduced bias to the analysis

## Conclusion

We propose a model in which spinal cord regeneration in zebrafish is supported by functionally distinct macrophage populations. Brain-derived microglia are recruited to the lesion, where they proliferate and differentiate into specialized states. Concurrently, CHT-derived BDMs infiltrate the injury site and exhibit substantial phenotypic plasticity, even acquiring microglial-like transcriptional programs. We identify at least seven distinct macrophage subpopulations with specialized phagocytic, proliferative, and inflammatory cell states. Importantly, we identify *g0s2* as a robust marker of microglia. Together, these findings provide the first detailed characterization of macrophage populations at the spinal cord injury site in zebrafish and establish a framework for understanding the complexity of immune cells in regeneration. Overall, our study suggests that the regenerative success of the zebrafish spinal cord is linked to a heterogeneous and dynamic response of microglia and BDM subpopulations.

## MATERIALS AND METHODS

### Ethics Statement

All experiments were performed under state of Saxony licenses TVV 36/2021, TVV 45/2018, and holding licenses DD24-5131/364/11, DD24-5131/364/12 (approved by Landesdirektion Sachsen, Referat 25 Veterinärwesen und Lebensmittelüberwachung); conformed to the guidelines established by the European Communities Council Directive of 22 September 2010 (2010/63/UE) and were in accordance with German animal law and regulations (Tierschutzgesetz TierSchG version 18 May 2006 with all modifications included thereafter until the date of publication of this manuscript).

### Animals

All zebrafish lines were raised and kept under standard conditions. Zebrafish experiments were performed under the experimental licenses TVV 36/2021, TVV 45/2018, and holding licenses DD24-5131/364/11, DD24-5131/364/12 from the Free State of Saxony. For experiments, larvae up to an age of 5 dpf of the following lines were used: Tg(*mpeg1.1:mCherry*)^gl23^, abbreviated as *mpeg1:mCherry* (Ellett et al., 2011); TgBAC(*p2ry12:p2ry12-GFP*)^hdb3Tg^, abbreviated as *p2ry12:GFP* (Sieger et al., 2012); Tg(*fli1:eGFP*)^y1^, abbreviated as *fli1:eGFP* (Roman et al., 2002)*; Tg(mpx:GFP)^uwm1^, abbreviated as mpx:GFP* (Renshaw et al., 2006); *csf1ra^j4e1/j4e1^ × csf1rb^+/re01^,* abbreviated as *csf1ra/b -/-* (Oosterhof et al., 2018).

### Spinal cord lesion

Lesions of larvae were performed as previously described (Ohnmacht et al., 2016). Briefly, 3 dpf zebrafish larvae were deeply anaesthetized in E3 medium (Nüsslein-Volhard & Dahm, 2002) containing 0,02 % of MS-222 (Sigma). Larvae were then transferred to a 4% agarose plate. After removal of excess water, larvae were positioned laterally. The entire spinal cord was transectioned using a 30G syringe needle at the level of the 15^th^ myotome, without injuring the notochord.

### Time-lapse microscopy

Zebrafish larvae were prepared for time lapse microscopy based on our established protocol (https://ibidi.com/img/cms/support/UP/UP12_Zebrafish_Co_Culture.pdf). Briefly, double transgenic zebrafish larvae (*p2ry12:eGFP*; *mpeg1:mCherry, or mpeg1:mCherry; fli1:eGFP*) were lesioned at 3 dpf and anaesthetized with MS-222 (160 mg/L in E3 medium). Low melting point agarose (0.5 %, Biozym Plaque Agarose - Low melting Agarose #840100) was prepared in E3 medium and allowed to cool down to 37 °C before adding MS-222 (final concentration 80 mg/L). Anaesthetized zebrafish larvae were transferred to the agarose and positioned into individual wells of the chambered coverslip. Microscopy was performed with a Dragonfly Spinning Disk microscope (Andor/Oxford Instruments, Belfast, UK) with a temperature-controlled stage (28 °C) using a 20x objective (20x/0.75 U Plan SApo, Air, DIC, OLYMPUS, Tokyo, Japan). GFP was excited using a 488 nm laser (*p2ry12:GFP*: 35% laser power, 2000 ms exposure time; *fli1:eGFP*: 20% laser power, 200 ms exposure time). mCherry was excited with a 561 nm laser (*mpeg1:mCherry*: 20% laser power, 200 ms exposure time).

We acquired z-stacks of 43 optical sections, with a step size of 1 µm for *p2ry12:eGFP; mpeg1:mCherry* and 76 optical sections, with a step size of 1 µm for *fli1:eGFP; mpeg1:mCherry*. The interval between single frames of the time-lapse movies was 6 minutes. Time-lapses were performed from 3 hpl until 30 hpl (*p2ry12:eGFP; mpeg1:mCherry*) or 2 hpl until 17 hpl (*fli1:eGFP; mpeg1:mCherry*).

### Cell counting in time lapse images

To analyse microglia migrating from the brain towards the spinal cord, a 500 µm wide was placed at the end of the brain stem. Cells were counted manually inside the window when entering into the window from rostral (brain) and leaving the window from caudal (spinal cord) along the whole time-lapse movie.

To analyse BDMs invading the injury site, we placed a 200 µm wide line below the injury site (middle of the notochord) and we counted each cell that crosses the line in either direction coming from the CHT. Cells were separated in cells moving towards the injury site, moving from the injury area back to the CHT, and cells moving immediately to and from the injury site along the entire time lapse movies. For cells migrating from peripheral tissues the line size was extended to 300 µm to ensure detection of most cells.

To analyse BDMs migrating through blood vessels, we counted all the *mpeg1:mCherry* positive cells inside of the *fli1:eGFP* structures along the entire time lapse that migrate form the CHT to the injury site.

To analyse cell heterogeneity in the injury site, a 200×75 µm window was placed in the center of the injury site and cells were counted manually.

### Seurat and Differentially Expressed Gene Analysis

The Seurat object of the scRNA seq dataset from (Cavone et al., 2021) was used to perform a macrophage sub-clustering. Clusters identified by enriched expression of *mpeg1.1* were subset and sub-clustering was performed by using the FindNeighbors (reduction = pca, dimensions = 20) and FindCluster (resolution = 0.35) functions in of the Seurat package in R (Butler et al., 2018; Hao et al., 2021, 2024; Satija et al., 2015; Stuart et al., 2019).

To identify marker genes, the FindAllMarkers function from the Seurat package (SeuratObject version 4.0) was used (default parameters). The “Wilcoxon rank sum test” with p_val_adj (p-value adjusted) < 0.1 and avg_log2FC >= 0.25 (average log2 fold change) was used for selecting significantly expressed marker genes. Genes were ordered based on their agvg_log2FC in descending order for each macrophage subcluster. Genes were then manually selected based on established/predicted functions from prior literature (non-annotated genes were excluded), and relative representative expression compared to other clusters (pct.1 > 0.2, pct.2 < 0.2).

### AUC-based marker selection

To quantify the discriminative power of each gene, we computed an Area Under the Curve (AUC) score. For each gene, expression values were retrieved across all cells together with the corresponding cluster identity. Cells belonging to the target cluster were encoded as positive, while all other cells encoded as negative. For each remaining gene, a receiver operating characteristic (ROC) curve was generated using the pROC package in R (Robin et al., 2011), and the AUC was extracted as a measure of how well gene expression distinguished the target cluster from non-target cells. AUC values ranged from 0 (non-discriminative) to 1 (perfectly discriminative). To facilitate interpretation, AUC values were binned into three qualitative categories representing marker strength: Low specificity (0–0.5), moderate specificity (0.5–0.7), high specificity (0.7–1). Each gene was assigned to one of these categories based on its computed AUC. A bar-chart summary of AUC values was generated using ggplot2 in R (Wickham, 2016).

### *In situ* hybridization chain reaction (HCR-FISH)

Larvae were collected in E3 medium and incubated with 0.003 % 1-phenyl 2-thiourea (PTU) at 6 hpf to suppress melanocyte development. Larvae were fixed overnight in 4 % paraformaldehyde (PFA) at 4 °C, and then transferred to 100 % Methanol (MeOH) at −20 °C overnight for permeabilization. Larvae were then rehydrated in a graded series of MeOH (25% decreases) at room temperature (RT) and subsequently washed in phosphate buffered saline with 0.1 % Tween (PBST). Thereafter, larvae were treated with 500 μL proteinase K (Roche) at a concentration of 30 μg/mL for 45 min at RT, followed by post-fixation in 4 % PFA for 20 min RT, and washes with PBST (3 x 5 min). Pre-hybridisation was performed using 500 μL of pre-warmed probe hybridisation buffer (Molecular Instruments, LA, USA) for 30 min at 37 °C. The probe sets, which were designed based on the NCBI sequence by Molecular Instruments (Choi et al., 2018), were prepared using 2 pmol of each probe set in 500 μL of pre-warmed hybridisation buffer. Then, buffer was replaced with the diluted probe sets and incubated for 16 h at 37 °C. Samples were washed 4 x 15 min with pre-warmed probe washing buffer (Molecular Instruments) at 37 °C followed by 2 x 5 min in 5x sodium chloride sodium citrate with 0.1 % Tween (SSCT) at RT. Larvae were then treated with 500 μL of room temperature equilibrated amplification buffer (Molecular Instruments, LA, USA) for 30 min at RT. Hairpin RNA preparation was performed following the manufacturer’s instructions. Samples were incubated with Hairpin RNAs for 16 h in the dark at RT. Finally, samples were washed for 3 x 30 min in 5 x SSCT at RT and immersed in 70 % glycerol. Samples were imaged with a Zeiss LSM980 confocal microscope.

### RNAscope combined with immunohistochemical labelling

RNAscope fluorescent in situ hybridization was performed using the RNAscope® Multiplex Fluorescent Reagent Kit v2 (Advanced Cell Diagnostics, Bio-Techne) and adapted for whole-mount zebrafish larvae following the manufacturer’s protocol with modifications based on Gross-Thebing et al. (2020). Larvae at 24 hpl were fixed in 4 % paraformaldehyde (PFA) in phosphate-buffered saline (PBS) for 40 minutes at room temperature (RT). Following fixation, larvae were washed three times for 5 minutes each in 0.1 % PBST), then dehydrated at RT through a graded MeOH) series (25% steps): (5 min each). Larvae were stored in 100 % MeOH at −20 °C overnight. For probe hybridization, larvae were rehydrated through a reverse MeOH series (25 % steps) and air-dried for 10 minutes at RT, ensuring a thin residual film of fluid remained to preserve tissue morphology. Larvae were then treated with two drops of Protease Plus solution (ACD Bio) and incubated for 1 hour at 40 °C in a water bath. Probes were warmed to 40 °C for 10 minutes prior to use. Hybridization was carried out by incubating larvae in 50 μL of probe solution mixed with probes in a 50:1:1 ratio overnight at 40 °C. The following day, samples were washed in 0.2× SSC with 0.01 % Tween-20 (SSC-Tween) at RT in three sequential steps (5, 10, 15 min). To preserve tissue integrity, larvae were post-fixed in 4 % PFA for 10 minutes at RT, followed by another round of SSC-Tween washes.

Signal amplification was performed via sequential incubation in amplification reagents (1-3), with SSC-Tween washes in between each step as previously described. For probe detection, the appropriate hybridization to HRP (Horseradish Peroxidase) step was selected based on the probe channel for 15 minutes at 40 °C, washed, and then incubated with TSA (Tyramide Signal Amplification) Vivid fluorophore (PerkinElmer) diluted 1:1500 in TSA buffer for 30 minutes at 40 °C. Fluorophores used were selected based on the imaging setup and experimental needs. HRP activity was blocked by incubation with HRP blocker for 15 minutes at 40 °C. SSC-Tween buffer and amplification reagents were prepared as per manufacturer instructions.

Following, immunolabelling for 4C4 was performed as usual. Briefly, samples were incubated overnight with primary antibody (mouse anti-4C4, 1:50, European Collection of Authenticated Cell Cultures, 7.4.C4) at 4°C. Incubations were performed in a blocking buffer solution containing 2 % BSA, 1 % DMSO, 1 % Triton and 10 % Normal donkey serum, in PBST. Afterwards, larvae were washed three times for 30 minutes each in PBST and incubated overnight with corresponding secondary antibody (1:200, Jackson Immuno). Finally, samples were washed three times for 30 minutes each in PBST and transferred to glycerol for mounting.

### Immune cell counting in *in situ* labelling

All images were analysed in FIJI (Schindelin et al., 2012). All images were oriented with rostral to the left, and dorsal at the top. A standardized window measuring 75 µm in height and 400 µm in width was centred over the injury site in each image, directly above the notochord, spanning the full mediolateral width of the spinal cord. Cells were counted manually inside the window along the Z-stack.

### Kernel Density Estimation (KDE) and spatial enrichment analysis

Kernel Density Estimation (KDE) (Silverman, 1998) and spatial enrichment analysis were performed in R. Cell coordinates were extracted from HCR manual cell counts in ImageJ. To ensure consistent spatial alignment across samples, coordinates were transformed relative to anatomical landmarks (dorso-ventral and rostro-caudal), with respect to the injury site centre. To further contextualize spatial patterns, the injury site was radially subdivided into three concentric zones (lesion core, lesion interphase, and peripheral intact tissue) defined based on KDE-derived density contours, providing a consistent reference framework across samples. Reference points marking the centre and edges of the injury site were manually defined to enable consistent spatial segmentation. Cell coordinates were combined into unified data frames for downstream spatial analysis and visualisation. Visualisation was performed using ggplot2, using the centre of the injury site of each individual fish as reference (x=0, y=0), to allow spatial comparison across markers and samples. To quantify the spatial distribution, two-dimensional kernel density estimates were used for each marker. For each marker, the probability density over a 200 x 50 µm grid spanning the observed spatial range was estimated. KDEs were computed using a Gaussian kernel via the kde2d function from the MASS R package (Venables & Ripley, 2002). The output KDE grid was used for downstream statistical enrichment analyses and visualization.

To determine whether observed marker^+^/transgene^+^ cells exhibit significant spatial clustering beyond random chance, we compared their spatial distributions to null model using bootstrapped KDEs (Manly & Navarro, 2020). To establish a null distribution, 100 bootstrapped KDEs were generated by randomly redistributing the same number of cells (n=100) uniformly within the global spatial bounds. Afterwards, the observed density at each grid cell was compared to the empirical distribution from the bootstraps and computed one-sided p-values. To assess spatial relationships between clusters, pairwise correlations were calculated between KDE-enriched areas. For each pair of markers, the corresponding KDE enrichment maps were flattened into one-dimensional vectors, and Pearson’s correlation coefficient (r) was computed across all grid cells. Correlation coefficients (r) were interpreted as follow: |r| ≥ 0.5 meaningful spatial relationship, r ≥ 0.5 adjacent localization, and r ≤ –0.5 indicating complementary or mutually exclusive spatial distributions. Correlations with |r| < 0.5 were interpreted as spatially independent.

### Data availability

The scRNa-seq data used in this study can be accessed at ArrayExpress accession no. E-MTAB-10379 (for Cavone et al. 2021, *mpeg1.1*:*GFP*-FACS-sorted) or NCBI GEO accession no. GSE276092 (for Docampo-Seara et al., 2026, non-FACS-sorted, all cells).

### Figure preparation

Images were adjusted for brightness, contrast and intensity in FIJI. Drawings were made using CorelDraw X8 (Corel/Alludo, Ottawa, Canada). Figure plates were prepared using CorelDraw X8 (Corel/Alludo, Ottawa, Canada) and Adobe Photoshop 2024 (Adobe, CA, USA). All plots and statistical analyses not otherwise described above were completed using GraphPad Prism 10 (GraphPad Software, Boston, USA). Error bars in figures indicate standard error of the mean (SEM).

## Acknowledgements

We thank Drs Hella Hartmann and Ruth Hans for imaging advice; and Dr. Judith Konantz, Marika Fischer, and Silvio Kunadt for fish care. This work was supported by the Light Microscopy Facility, the DRESDEN-concept Genome Center, the Flow Cytometry Facility, and the Zebrafish Facility, all core facilities of the Center for Molecular and Cellular Bioengeneering (CMCB) at the Technische Universität (TU) Dresden. Funding was provided by an Alexander von Humboldt Stiftung Professorship award (to CGB) and TU Dresden core funding (to CGB).

## Author contributions

**KH**: conceptualization, methodology, validation, formal analysis, investigation, writing - original draft; **EJ:** investigation; **OC:** investigation; **SJE:** investigation, writing - original draft; **BW**: investigation; **CGB:** conceptualization, writing - original draft, writing - review and editing, supervision, funding acquisition; **TB:** conceptualization, writing - original draft, writing - review and editing, supervision, funding acquisition; **ADS:** conceptualization, methodology, validation, formal analysis, investigation, writing - original draft, writing - review and editing, visualisation, supervision.

## SUPPLEMENTARY FIGURES

**Suppl. Fig. 1.**
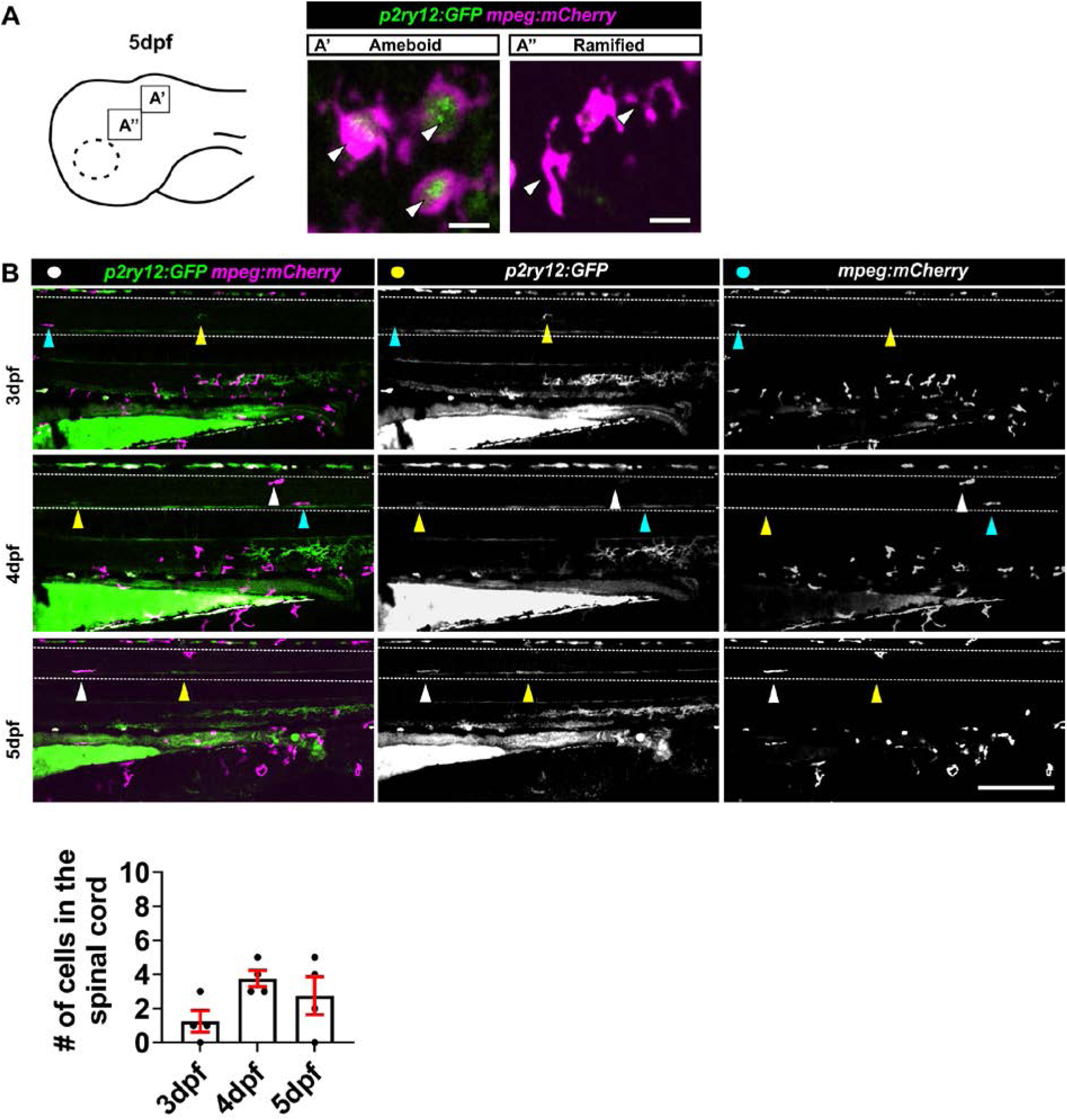
Analysis of microglia in unlesioned fish. **(A)** Photomicrograph of the head of a 5 dpf larvae showing expression of *p2ry12:GFP* and *mpeg1:mCherry*. **(A’-A’’’)** High magnification images showing amoeboid and ramified like microglia (arrows). Note that *p2ry12* expression is low or below detection threshold. **(B)** Images and quantification at 3, 4, and 5 dpf showing that few cells are present within the spinal cord (1.3 ± 0.63 at 3 dpf, 3.7 ± 0.48 at 4 dpf, 2.8 ± 1.11 at 5 dpf). White arrows label double-positive cells, yellow arrows *p2ry12:GFP*-only positive cells, and blue arrows label *mpeg1:mCherry-*positive cells with low or minimal *p2ry12:GFP* signal. Dotted lines delineate the spinal cord. **Scale bars:** 200 µm (A), 100 µm (B), 15 µm (A’-A’’’).

**Suppl. Fig. 2.**
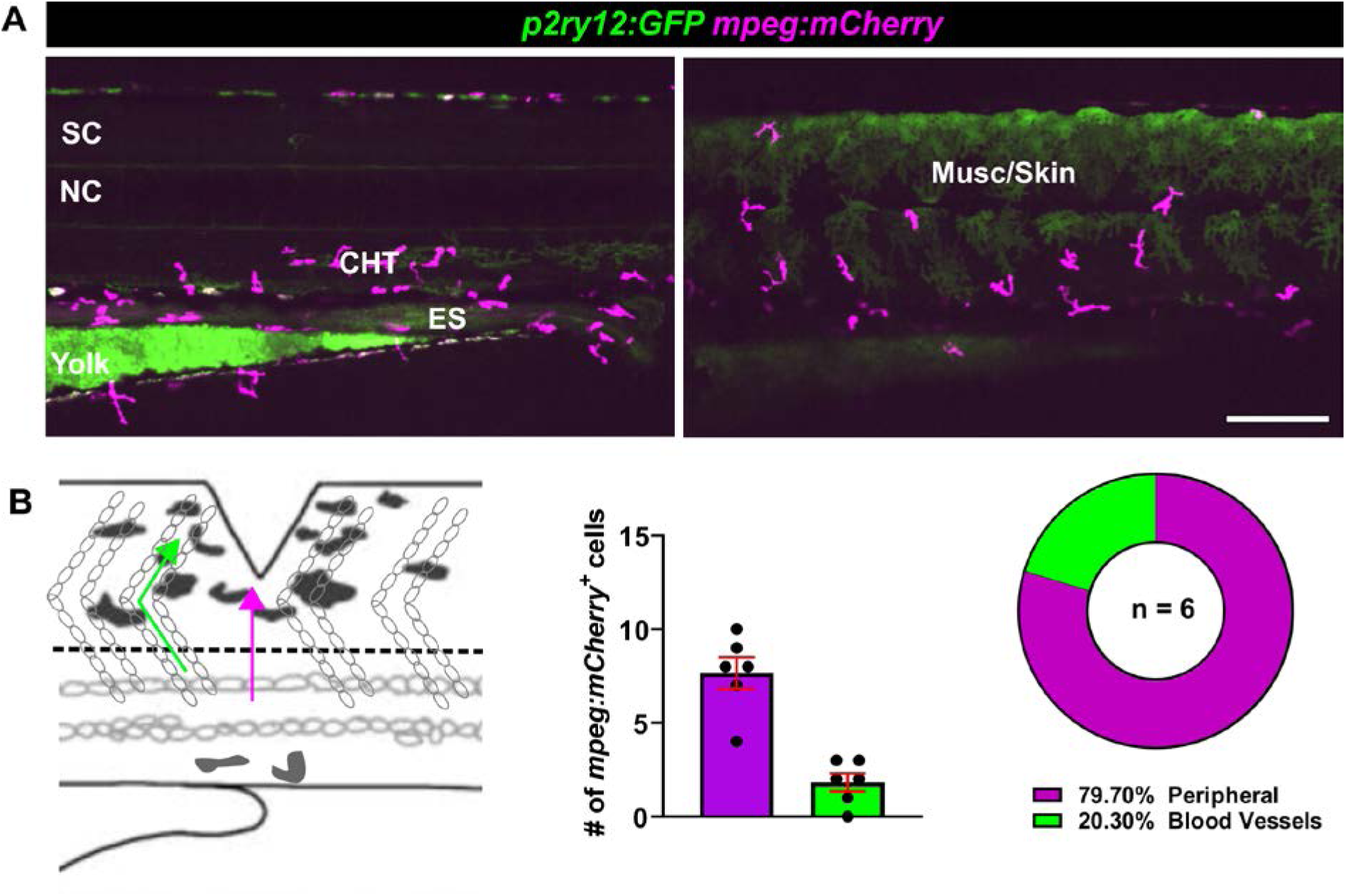
BDMs reach the lesion through tissue migration and via blood vessels. **(A)** Images showing presence of *mpeg1:mCherry* positive cells in unlesioned fish in the CHT, enteric system and muscle/skin tissue. **(B)** Quantification of time-lapse movies for *mpeg1:mCherry* / *fli1:eGFP* transgenic animals, showing numbers and percentages of cells migrating to the injury site from the CHT via peripheral tissue (magenta) or thought the blood vessels (green). Few *mpeg1:mCherry* positive cells migrate via blood vessels (peripheral: 7.7 ± 0.84; blood-vessels: 1.83 ± 0.48). **Scale bars:** 100 µm.

**Suppl. Fig. 3.**
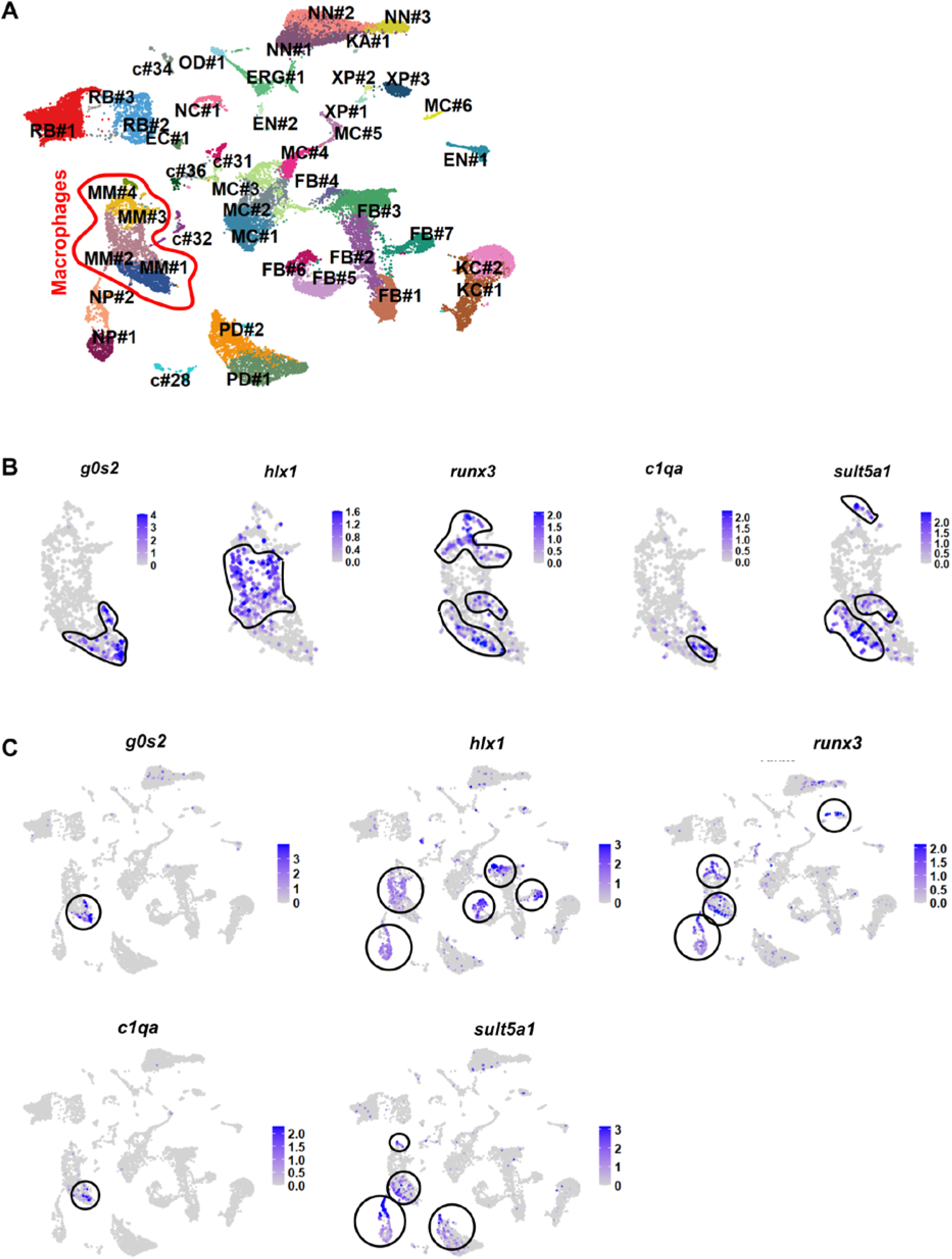
Analysis of selected marker genes in a non-FACS-sorted scRNA-seq dataset. **(A)** UMAP from non-FACS-sorted scRNA-seq dataset from Docampo-Seara et al. 2026 at 24 hpl showing all cells in the injury site. **(B)** Feature plots of marker expression in macrophages. Contours define positive cell areas. **(C)** Feature plots of selected genes in the entire dataset showing that *g0s2* and *c1qa* show high macrophage expression specificity, while *hlx1, runx3* and *sult5a1* are also expressed in other cell clusters (circles highlight the expression). Abbreviations: MM (macrophages), NP (neutrophils), PD (peridermis), FB (fibroblasts), KC (keratinocytes), MC (muscle cells), XP (xanotphores), MC (melanocytes), EN (enteric nervous system), NN (neurons), KA (Kolmer-Agduhr Cells), ERG (ependymo-radial glial cells), NC (notochord), OD (oligodendrocytes), RB (red-blood cells), c (unknown).

**Suppl. Fig. 4.**
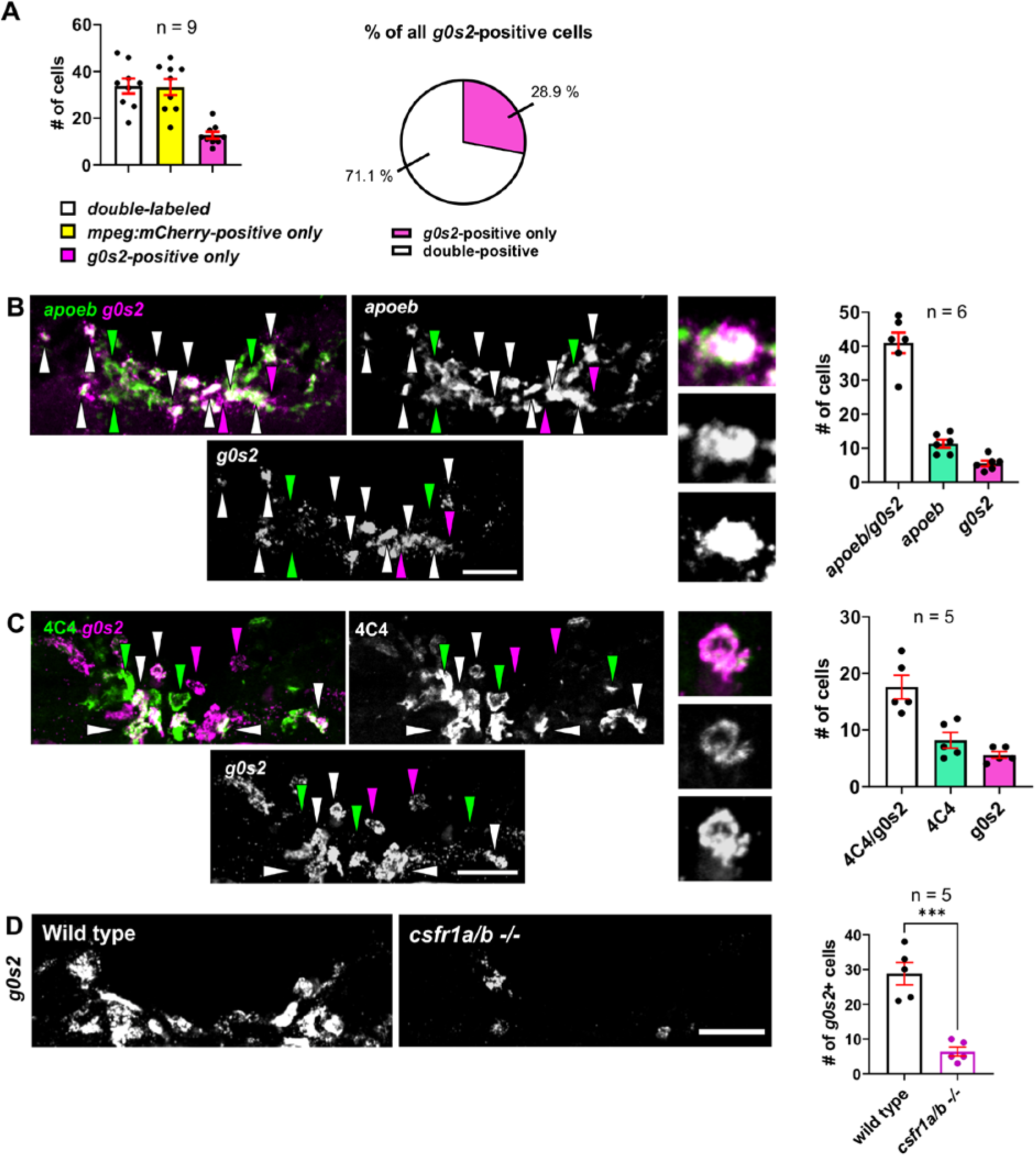
*g0s2* is a putative microglia marker. **(A)** All cell quantification of HCR for *g0s2* in *mpeg1:mCherry* line (performed in Fig. 5A) showing that ∼29% out of all *g0s2*-positive cells are negative for *mpeg1:mCherry* (double-positive cells: 33.78 ± 3.2cells; *mpeg1:mCherry*-only: 33.33 ± 3.4cells; *g0s2*-only: 12.78 ± 1.4cells) **(B)** HCR-FISH labelling of *apoeb* and *g0s2* is shown. Green arrows point at *apoeb* only positive cells, pink arrows point at *g0s2*-only positive cells and white arrows point at double positive cells. Of all *g0s2*-positive cells, 88% were also *apoeb*-positive. **(C)** RNAscope and immunohistochemical labelling of 4C4 and *g0s2* showing colocalization between both markers. Green arrows point at 4C4 single positive cells, pink arrows point at *g0s2* single positive cells and white arrows point at double positive cells. Of all *g0s2*-positive cells, 75% were 4C4-positive. **(D)** RNAscope analysis of *g0s2* in wild type and *csfr1a/b* double -/- fish, shows an 80% lower number of *g0s2* positive cells in the mutant compared to wild type (wild type: 28.8 ± 3.2cells per larvae; *csfr1a/b*-/-: 6.4 ± 1.3cells per larvae; Unpaired t-test: 0.0002). **Scale bars:** 50 µm.

**Suppl. Fig. 5.**
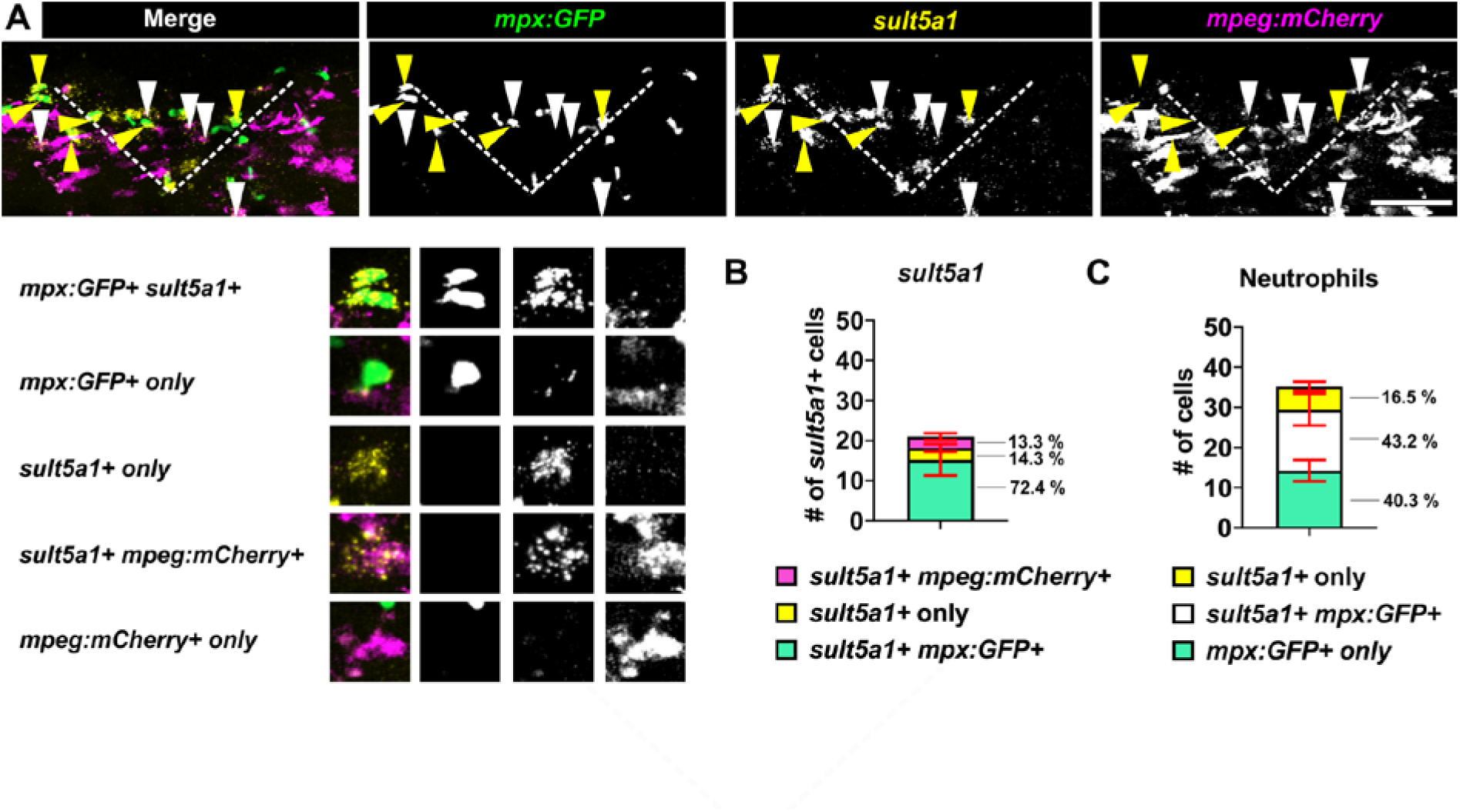
*sult5a1* is also expressed in a cluster of neutrophils. **(A)** HCR in *mpx:GFP/mpeg1:mCherry* double transgenic line at 24hpl showing *sult5a1* expression in neutrophils (yellow arrows) and in macrophages (white arrows) **(B)** Quantification of the number of all *sult5a1*-positive cells showing that the vast majority of cells (72.4%) are neutrophils. **(C)** Quantification showing that almost half of the neutrophils are *sult5a1*-positive. **Scale bars**: 50 µm.

**Suppl. Fig. 6.**
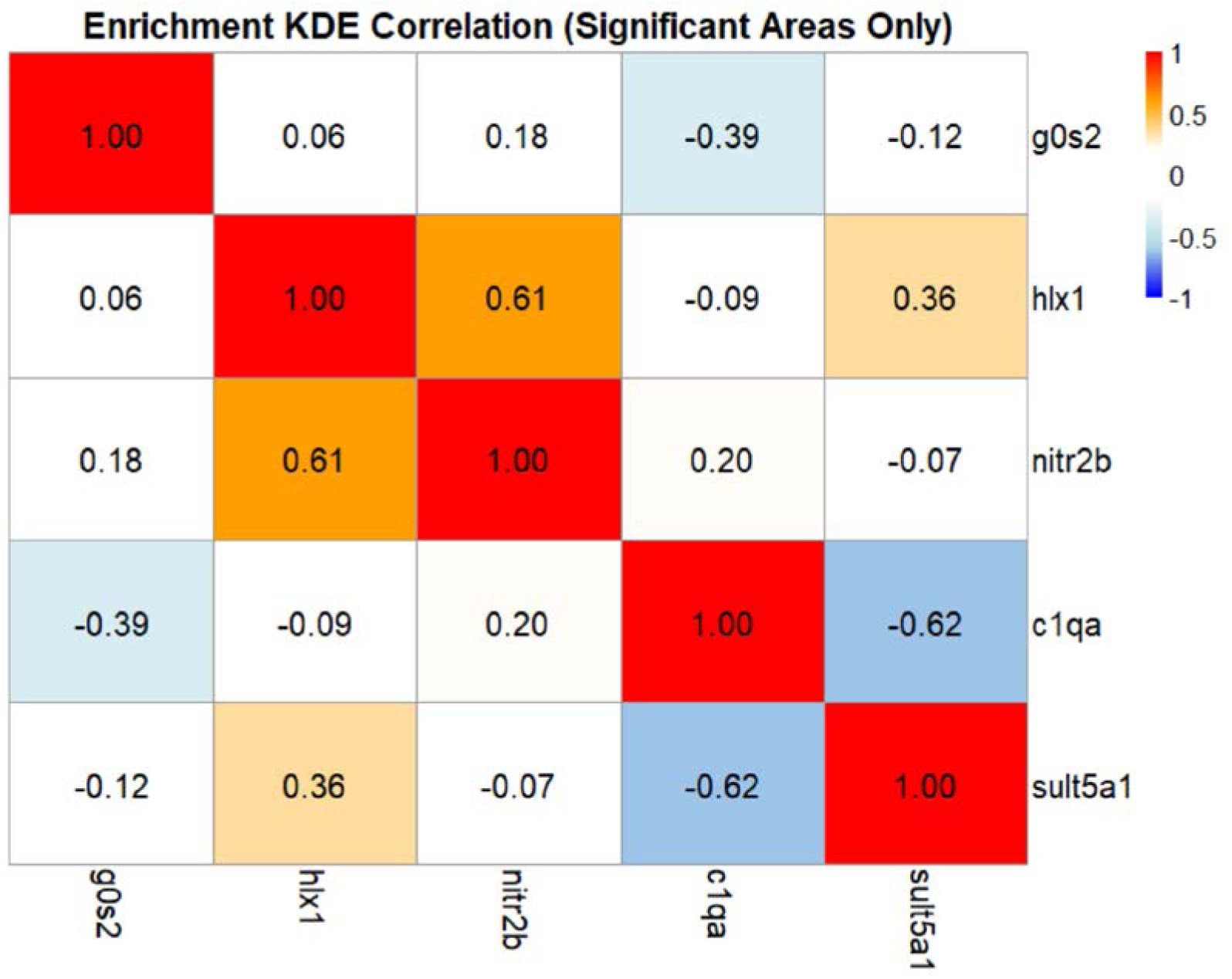
Correlation matrix indicating cluster spatial correlation. 1 indicates a total correlation, 0.5 moderate, 0 no correlation, and −1 total negative correlation. Absolute correlation is shown to each cluster compared to itself. Results show that clusters do not show a spatial correlation, with the exception of cluster 2 (*hlx1*) and cluster 4 (*nitr2b*), that showed a moderate correlation (between 0 and 0.5).

## Supplemental Data File Legends

**S1 movie:** Time lapse movie from 3 hpl to 33 hpl using the *mpeg1:mCherry* and *P2ry12:GFP* reporter lines.

**S2 movie:** Time lapse movie from 2 hpl to 19 hpl using the *mpeg1:mCherry* and *fli1:eGFP* reporter lines.

